# Quantitative Dynamic Analysis of *de novo* Sphingolipid Biosynthesis in *Arabidopsis thaliana*

**DOI:** 10.1101/2023.12.08.570827

**Authors:** Abraham Osinuga, Ariadna Gonzalez Solis, Rebecca E. Cahoon, Adil Al-Siyabi, Edgar B. Cahoon, Rajib Saha

## Abstract

Sphingolipids are pivotal for plant development and stress responses. Growing interest has been directed towards fully comprehending the regulatory mechanisms of the sphingolipid pathway. We explore its *de novo* biosynthesis and homeostasis in *Arabidopsis thaliana* cell cultures, shedding light on fundamental metabolic mechanisms. Employing ^15^N isotope labeling and quantitative dynamic modeling approach, we developed a **r**egularized and constraint-based **D**ynamic **M**etabolic **F**lux **A**nalysis (r-DMFA) framework to predict metabolic shifts due to enzymatic changes. Our analysis revealed key enzymes such as sphingoid-base hydroxylase (SBH) and long-chain-base kinase (LCBK) to be critical for maintaining sphingolipid homeostasis. Disruptions in these enzymes were found to affect cellular viability and increase the potential for programmed cell death (PCD). Thus, this work enhances our understanding of sphingolipid metabolism and demonstrates the utility of dynamic modeling in analyzing complex metabolic pathways.

## Introduction

Sphingolipids represent a diverse class of lipids that mesh within the cellular structure of eukaryotic cells. Their significance is underscored by their dual role: serving as abundant component of cell membranes and bioactive molecules like ceramide (Cer) and sphingoid bases (Michaelson et al., 2016; Snider et al., 2018). With plants, typically, the significance of sphingolipids is undeniable. Constituting up to 40% of the plasma membrane, these lipids prominently feature in cellular endomembranes, such as the endoplasmic reticulum (ER), golgi, and tonoplast (Buré et al., 2014; Luttgeharm et al., 2016). Owing to their structural uniqueness, they have essential cellular functions. For instance, glycosylated variants such as glucosylceramides (GlcCer) and glycosylinositolphosphoceramides (GIPCs) not only bolster membrane function but also facilitate protein trafficking in the cell (Michaelson et al., 2016). Moreover, under environmental stress, such as microbial pathogenesis, an accumulation of specific sphingolipids triggers the onset of programmed cell death (PCD) (Magnin-Robert et al., 2015; Worrall et al., 2008).

With these molecules crucially situated in cell membranes and influencing cell signaling pathways (Merrill, 2002), the sphingolipid biosynthesis (*de novo)*, initiated in the endoplasmic reticulum (ER), is central to cellular function and homeostasis (Merrill, 2002, 2011). The pathway specific to *Arabidopsis thaliana* (hereafter Arabidopsis), visualized later in Fig. 2A, commences with serine palmitoyltransferase (SPT) catalyzing a foundational reaction involving palmitoyl-CoA and serine to produce 3-ketosphinganine, a rate-limiting phase in sphingolipids backbone, sphingoid or long-chain base (LCB), synthesis (Hanada, 2003). The ensuing enzymatic processes yield a variety of ceramides, further differentiated by ceramide synthases in Arabidopsis tailored to distinct acyl-CoA lengths (Markham et al., 2011, 2013; Ternes et al., 2011). These ceramides, through glycosylation, form glucosylceramides and further steps contribute to the synthesis of GIPCs which are dominant glycosphingolipids in plant cells (Luttgeharm et al., 2016; Msanne et al., 2015). The limiting and tight regulation, led by SPT (Hanada, 2003), ensures balanced sphingolipid production while avoiding undue aggregation of PCD-inducing components.

In plant biology, the significance of sphingolipid biosynthesis is underscored by its regulatory responses to diverse stimuli, encompassing both biotic interactions and abiotic environmental factors. The interplay of genes, enzymes, and signaling molecules is fundamental to maintaining cellular homeostasis and facilitating adaptability to internal or external perturbations (Greenberg et al., 2011; Nair et al., 2019). Depending on the nature of perturbations—whether pathogenic or otherwise—cellular responses can manifest as transient dynamics or prolonged steady-state adjustments, reflecting the metabolic pathway’s adaptability across diverse physiological conditions. Notably, in plants, yeast and human cells, the phosphorylation of sphingolipids and associated enzymes emerges as a pivotal, albeit not entirely deciphered, regulatory process (Guo & Wang, 2012; Imai & Nishiura, 2005; Qin et al., 2017; Worrall et al., 2008). Studies suggest a connection between these phosphorylation events and signaling molecules, potentially triggering the production of reactive oxidative species, which are in turn implicated in programmed cell death (PCD) pathways (Guo & Wang, 2012; Ray et al., 2012; Shi et al., 2007). Because of this, enzymes such as long-chain-base kinase (LCBK) and sphingolipid-base hydroxylase (SBH) are noteworthy, as well as a putative sphingoid-phosphate phosphatase (LCBPP1) which is believed to dephosphorylate sphingoid molecules restoring them to their long-chain base precursor forms. Thus, these enzymes emerge as potential regulators in response to an external stimulus/stress by modulating the balance of phosphorylated/dephosphorylated species of the sphingolipid biosynthesis pathway (Aguilera-Romero et al., 2014). These enzymes are particularly intriguing due to their evident association with the aforementioned processes. The specific dynamics or interactions of *de novo* synthesis path that might involve these kinases, or how the enzymes’ functions might impact broader cellular or physiological responses, has not been looked into. However, both free LCBs and phosphorylated forms typically occur in low abundance in plants making it difficult to measure them (Markham & Jaworski, 2007).

To elucidate the sphingolipid pathway, stable isotope labeling has become an invaluable technique (Ecker & Liebisch, 2014; Wigger et al., 2019), leveraging tools such as mass spectrometry (MS) and nuclear magnetic resonance spectroscopy (NMR) (Allen & Young, 2020). Although ^13^C remains the popular choice for labeling, ^15^N and ^18^O isotopes are gaining momentum (Allen & Young, 2020). In autotrophs, isotopically labeled nutrients, often introduced as nitrate salts, provide invaluable tracers for monitoring metabolic flows under real-life conditions (Kuchař et al., 2016; S. Li et al., 2013; Martínez-Montañés & Schneiter, 2016). This method’s efficacy is highlighted in its application to organisms like *Saccharomyces cerevisiae* (baker’s yeast) and Arabidopsis, unearthing insights into lipid and photosynthetic processes. However, with sphingolipids, given their essential role in various cellular processes, accurate detection and quantification is necessary. Owing to their unparalleled sensitivities and specificities, MS tandem MS (MS/MS) based approach has become a gold standard for detection and quantification tasks, ensuring precise measurements of these vital molecules in a variety of contexts. This analytical method’s precision not only allows for the clear identification of sphingolipids but also enables detailed insights into their intricate structures and molecular dynamics (Allen et al., 2015; Ecker & Liebisch, 2014; Kuchař et al., 2016; Martínez-Montañés & Schneiter, 2016).

In the multifaceted realm of sphingolipid metabolism and the dynamic nature of plant metabolism, tools for accurate exploration of these pathways are vital. Computational systems biology addresses this, either affirming established hypotheses or generating new ones from observed patterns. In recent times, constraint-based genome-scale metabolic models (cGMMs) have been crafted to probe the fundamental metabolism of several plant models, with *Arabidopsis* being a notable subject (Schroeder & Saha, 2020). Isotopic labeling-based Metabolic Flux Analysis (MFA) offers a quantitative, mechanistic portrayal of cellular phenotypes, producing a flux map grounded on the stoichiometric balance of isotopic labeling patterns (Antoniewicz, 2013a). Typical steady state MFA offers an initial glimpse into labeled product distributions, but it doesn’t directly discern intermediate turnover rates. This limitation is addressed by employing a detailed non-stationary isotope dynamic metabolic profiling, which mechanistically yields turnover rates and transient metabolic fluxes (Allen & Young, 2020; L. Li et al., 2017; Ma et al., 2014; Wall et al., 2015). A pioneering technique in this area is Dynamic Metabolic Flux Analysis (DMFA), which can be viewed as an enhancement of MFA (Antoniewicz, 2013b), specifically, isotope tracer experiments incorporating time-dependent features for an exhaustive biological analysis.

DMFA consists of two primary strategies: the Static Optimization-Based Approach (SOA) and the Dynamic Optimization-Based Approach (DOA) as outlined by Antoniewicz (Antoniewicz, 2013b). SOAs simply average data, segmenting culture time into metabolic phases, or use data smoothing techniques for a stepwise solution approach. DOAs, on the other hand, use piecewise linear flux functions across continuous time segments. While comprehensive dynamic optimization would be ideal for accuracy, it’s often impractical for extensive models. Conversely, SOAs may compromise on accuracy and solution stability. However, SOA has been employed in various biological organisms modeling, from *Escherichia coli* (E. coli) to Arabidopsis. DOA, with its distinct benefits, has also been used for *E. coli* metabolism and signaling pathways in S. cerevisiae (Antoniewicz, 2013a; Hackett et al., 2016). As exploration in this domain widens, these strategies have set the stage for groundbreaking studies on plant metabolism, like resource allocation in Arabidopsis. Nonetheless, DMFA’s effectiveness is tethered to technological advancements. With the emergence of high-throughput metabolomics and computational modeling, dynamic models of metabolism (dMMs) have been established (Di Filippo et al., 2022). Similar to DMFA, dMMs are also generally crucial for grasping the metabolic regulation of cellular perturbations over time durations. However, predicting the repercussions of such disruptions via dMMs remains crucial but challenging, considering the extensive regulatory and feedback mechanisms involved.

Considering the regulations and broad impacts of sphingolipid metabolism in diverse biological contexts, our study conducts a quantitative dynamic metabolic flux analysis on the *de novo* sphingolipid biosynthesis in Arabidopsis. While numerous investigations have delved into the regulation of the *de novo* sphingolipid biosynthesis, employing tools like isotope dynamic labeling in yeast and mammalian cells, the detailed dynamic metabolic flux analysis of *de novo* synthesis of sphingolipids in Arabidopsis remains relatively unexplored (Haynes et al., 2011; Siow et al., 2015; Snider et al., 2018; Wigger et al., 2019; You et al., 2014). As an example, Chen, Alonso, and Shachar-Hill (2013) (X. Chen et al., 2013) developed a model rooted in pulse labeling, but it didn’t focus on *de novo* synthesis. This line of inquiry is essential given the potential hazards of alterations in *de novo* sphingolipid biosynthesis. Well-documented cases, such as the effects of fumonisins—a mycotoxin found in maize—highlight these hazards, with outcomes ranging from diseases in plants to cancer in mammals. Within the realm of sphingolipid metabolism, utilizing DMFA to track temporal changes offers insights into metabolic stability, which is crucial for therapeutic explorations. A nuanced analysis of metabolic flux shifts can pinpoint specific molecular subspecies or enzymes, enabling targeted modifications to regulate downstream processes.

In this work, we used a time-course ^15^N isotopic labeling technique coupled with targeted metabolomics to gain a dynamic and quantitative understanding of the *de novo* sphingolipid biosynthesis in T87 cell cultures. We delved deeper using a DMFA framework, which incorporated both regularization and dynamic flux sampling, termed r-DMFA. This approach unveiled temporal variations associated with enzymatic changes. Fig. 1 provides a comprehensive view of our study workflow and methodology. Our computational analysis emphasized the pivotal role of SBH in preserving cellular equilibrium. A shortage of SBH led to distinct metabolic shifts and an increased predisposition to PCD, a consequence of sphinganine accumulation. Notably, active long-chain-base kinase (LCBK) channel, which comprises of its analogous and putative reversible enzyme, was identified to regulate shortage SBH activity and by potentially acting as a rate-determining factor in the degradation of phosphorylated phytosphinganine/phytosphingosine. Additionally, our in-depth flux analysis also showcased a cooperative relationship among FA2H, CSII, and GCS enzymes—key pillars of cellular stability. Disturbances in their functions jeopardized cell viability and raised PCD risks, as evident from potential LCB/Cer build-up and deviations from standard wild-type (WT) dynamics. From these insights, we suggest particular control nodes in the sphingolipid synthesis pathway. Our findings aim to create a predictive model of the sphingolipid network’s response to genetic and environmental changes, setting the stage for bioengineering stress-resilient crops.

**Fig. 1.**
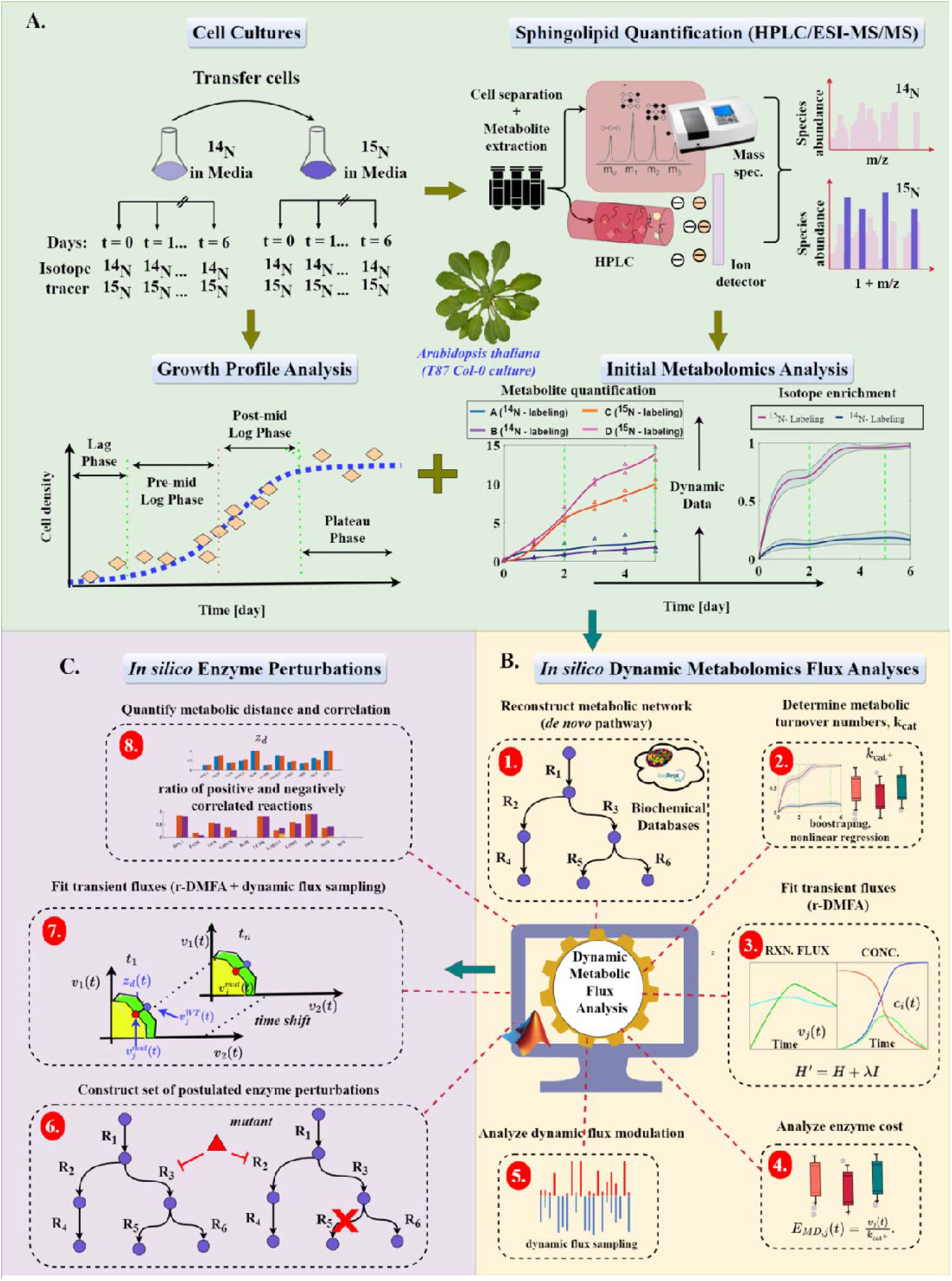
Overview of the workflow and methodology used in this work. **(A)** Experiment design and labeling workflow depicting cell culture, growth, isotope labeling, quantification, and initial metabolomics analysis. (B) Schematic illustration of the step-by-step procedure employed to obtain transient metabolic fluxes and analyze the inherent dynamic modulations of the Arabidopsis sphingolipid biosynthesis pathway. (C) *In silico* knockout (KO)/essentiality hypothesis testing using the customized MATLAB implementation of r-DMFA modeling framework. Within the metabolic network, hypothetical enzyme perturbations were introduced and assessed using the r-DMFA modeling framework in MATLAB, highlighting transient flux alterations. By comparing the model-generated fluxes with those observed in a wild-type reference model, this method efficiently guides and streamlines subsequent experimental validations, reducing the need for repeated iterations.

## Results and Discussion

### Isotopic dynamic labeling and sphingolipids quantification

In the pursuit of a comprehensive method to quantify diverse sphingolipids in raw samples, mass spectrometry (MS) was chosen for its adaptability to individual analytes. Due to the complexity of plant sphingolipid structures, MS was combined with high-performance liquid chromatography (HPLC). This combination facilitated efficient transient sphingolipid measurements and conformed to the principles of the HPLC/electrospray ionization tandem mass spectrometry (HPLC/ESI-MS/MS) method as detailed in (Cahoon et al., 2021; Markham & Jaworski, 2007). For quantifying sphingolipids, a prevailing challenge involved in ^13^C-labeling via MS, as prior studies typically reveal are issues in transitions, modifications, and fragmentation (Snider et al., 2018). To address this, ^15^N stable isotope labeling was utilized, targeting the nitrogen atom in the long-chain base, to provide a direct quantification approach for various sphingolipid species. Each molecule of LCB, ceramide (Cer), hydroxyceramide (hCer), and glucosylceramide (GlcCer) contains one nitrogen-15 atom due to the single nitrogen in their long-chain base. Although Arabidopsis glycosylinositol phosphorylceramides (GIPCs) may occasionally feature a second nitrogen atom on one of their sugar residues, this modification is uncommon in Arabidopsis leaves and its frequency in cultured cells is unknown. The presence of a second ^15^N in certain GIPC varieties does not impact these results, but it should be noted.

In an *in vitro* experiment, unlabeled *A. thaliana* T87 cell suspension cultures established from the ecotype Columbia, were grown in NT-1 liquid medium with the naturally occurring ^14^N isotope. ^15^N–labeled cultures were fed with NT-1 liquid medium containing K^15^NO_3_ and ^15^NH_4_ ^15^NO_3_ (Cambridge Isotope Labs) to introduce a stable ^15^N isotope under the same conditions. HPLC/ESI-MS/MS was used to quantify 168 sphingolipid species over a time course. The incorporation of ^15^N into sphingolipids was monitored as a +1 m/z mass increase for both precursor and product ions for multiple reaction monitoring (MRM) transitions.

Following HPLC/ESI-MS/MS analysis, sphingolipid molecular species were measured until the mid-exponential growth phase, which occurs on Day 6 (see Supplementary Fig. 1). Fig. 2B displays the total detected sphingolipids and highlights the abundance of the ^14^N and ^15^N isotopes for each species on Day 5. This day is representative of the mid-exponential growth phase. The fingerprint patterns observed for each sphingolipid class in Fig. 2B are consistent with the findings of Markham and Jaworski (Markham & Jaworski, 2007) for crude samples of Arabidopsis thaliana Col-0. Moreover, due to sphingolipids’ essential roles in plant cellular processes like growth, differentiation, and stress response, particularly in Arabidopsis, we deliberately focused on what happened up to the mid-exponential growth phase (Luttgeharm et al., 2016; Markham et al., 2013). The mid exponential growth phase is marked by peak cell division and growth, and it has significant physiological implications. During this phase, there’s an increased demand for membrane components, which can lead to notable changes in sphingolipid metabolism. By narrowing the focus to this phase, potential complications from later stages of growth were circumvented. As a result, a foundational dataset was obtained, which will be beneficial for further studies and future research.

**Fig. 2.**
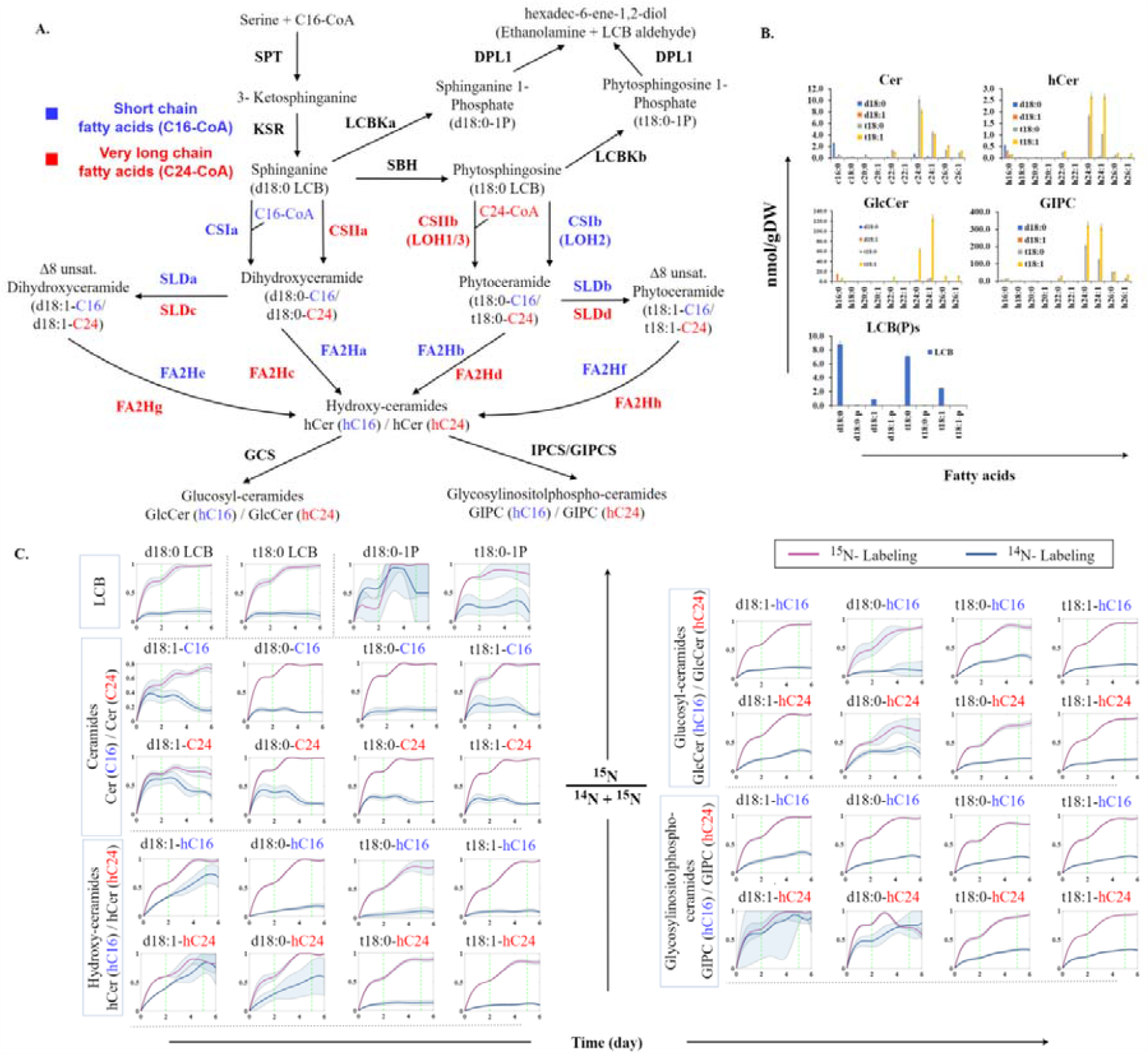
Comprehensive ^15^N profiling and isotopologue analysis of sphingolipid species. **(A)** Overview of the sphingolipid biosynthesis pathway. The illustrated pathway model, tailored for r-DMFA analysis, aligns with the model detailed by (Alsiyabi et al., 2021). This representation captures the full spectrum of active metabolic transformations in Arabidopsis sphingolipid biosynthesis. It integrates specific enzyme activities and streamlines metabolic pathway steps, especially where existing literature is limited. The model accommodates enzyme promiscuity, underscores minimal enzyme activity, and acknowledges the dominance of either C16 or C24 fatty acids (FA) in sphingolipids. We’ve also excluded certain reactions, notably those converting non-hydroxylated ceramides to complex forms, consistent with the primary composition of glycosylated sphingolipids. For clarity, blue elements relate to C16 fatty acids, while red pertains to C24, with accompanying enzyme annotations (letters a-h) for specificity. SPT: serine palmitoyltransferase, KSR: 3-ketosphinganine reductase, SBH: LCB C-4 hydroxylase, CSI: class I ceramide synthase, CSII: class II ceramide synthase, SLD: LCB Δ8 desaturase, FA2H: fatty acid hydroxylase, GCS: glucosylceramide synthase, GlcCer: glucosylceramide, GIPCS: glycosyl inositolphosphoceramide synthase, LCB: long chain base, LCBK: long chain base kinase and DPL1: dihydrosphinganine phosphate lyase. Letters a-h indicate the enzyme reaction specificity to the leading reactive species. **(B)** A quantitative snapshot of ^15^N-labeled sphingolipids in Arabidopsis T87 cell cultures. This captures the isotopic abundance of sphingolipids on day 5 of the growth phase, with patterns resonating with those in Arabidopsis Col-0 crude samples as reported by Markham & Jaworski (2007). We’ve provided distinct notations to differentiate between saturated, unsaturated, and hydroxylated forms of both dihydroxy and trihydroxy LCB groups and FA entities. For a more granular view of the growth trajectory, see Supplementary Fig. 1. Saturated forms of dihydroxy and trihydroxy LCB groups are represented by “d18:0”, “t18:0” while the unsaturated forms are represented as “d18:1” and “t18:1”, respectively. FA groups in saturated and unsaturated forms for example with 16 carbon chain, C16 are represented as “c16:0”, “c16:1” while the hydroxylated forms are “h16:0”, “h16:1”. **(C)** Transient isotopologue fractions [(m + 1)/(m + 1 + m + 0)] of 36 sphingolipid species during ^15^N labeling. Each depicted fraction represents the proportion of ^15^N enrichment in the measured analyte relative to its entire pool. The labeling also accounts for both pre-existing analytes and those integrated with ^15^N during the experiment. The patterns observed highlight the adaptability of the introduced ^15^N label in different sphingolipid species, with specific attention to their rate of incorporation over time.

To dynamically trace ^15^N incorporation into sphingolipid species, isotopologues, base m/z (m + 0) and +1 m/z shift (m + 1), of labeled and unlabeled sphingolipid species (hereafter termed ‘analyte’) were analyzed. The (m + 0) isotopologue, exemplified by 302.3 m/z for “d18:0”, represents the molecule’s mass where all atoms exist in their predominant isotopic form, notably ^14^N. In contrast, the (m + 1) isotopologue, such as 303.3 m/z for “d18:0”, denotes the mass shift resulting from ^15^N integration or from the natural presence of ^13^C, ^18^O, and ^2^H isotopes. The similar trends observed in the entire active pool measurements (m + 1 + m + 0) across biological replicates under both labeled and unlabeled conditions (as depicted in Supplementary Fig. 2 and supported by Supplementary Fig. 3) attest to effectiveness of this approach. Through dynamic ^15^N-labeling, isotopologue fractions [(m + 1)/(m + 1 + m + 0)] were computed, which signify the proportion of ^15^N enrichment in the analyte relative to its entire pool. As such, isotopologue fractions during ^15^N labeling are internally adjusted to account for the quantity of pre-existing analytes prior to ^15^N integration, as depicted in Fig. 2C.

Fig. 2C further illustrates transient isotopologue fractions of 36 sphingolipid species. A ^14^N-labeled control was employed for each species to assert measurement specificity. This measure is anticipated to remain consistent. Corresponding temporal ^15^N abundances, indicative of *de-novo* synthesis, are portrayed in Supplementary Fig. 3. The LCB class of sphingolipids, instrumental due to their association with rate-limiting steps of this pathway, was notably abundant. Meanwhile, temporal isotopologue fractions of LCB phosphorylated forms (LCBPs) exhibited variances which could be attributed to their low abundance, inactive pools, or instrument sensitivity. In labeled sphingolipids with trihydroxy LCB backbones (“t18:0”/ “t18:1”), an increase in (m + 1, for ^15^N) and a corresponding decrease in (m + 0, for ^14^N) ion counts were observed. These patterns, especially evident during the early log-growth phase between days 2 and 5 of ^15^N labeling, mirror trends witnessed in dihydroxy LCB derivative sphingolipids (“d18:0”/ “d18:1”). Nonetheless, the (m + 1) fractions of unlabeled sphingolipids remained mostly invariant throughout the labeling interval, as evidenced in Fig. 2C and 3. As represented in Fig. 2C and 3, ^15^N enrichment in intracellular sphingolipids reached an 80-100% plateau by day 5, sustaining thereafter. This alteration in the “heavy” (m + 1) fraction relative to the unlabeled condition underscores the utility of nitrogen isotopes for dynamic labeling in newly synthesized sphingolipid species.

**Fig. 3.**
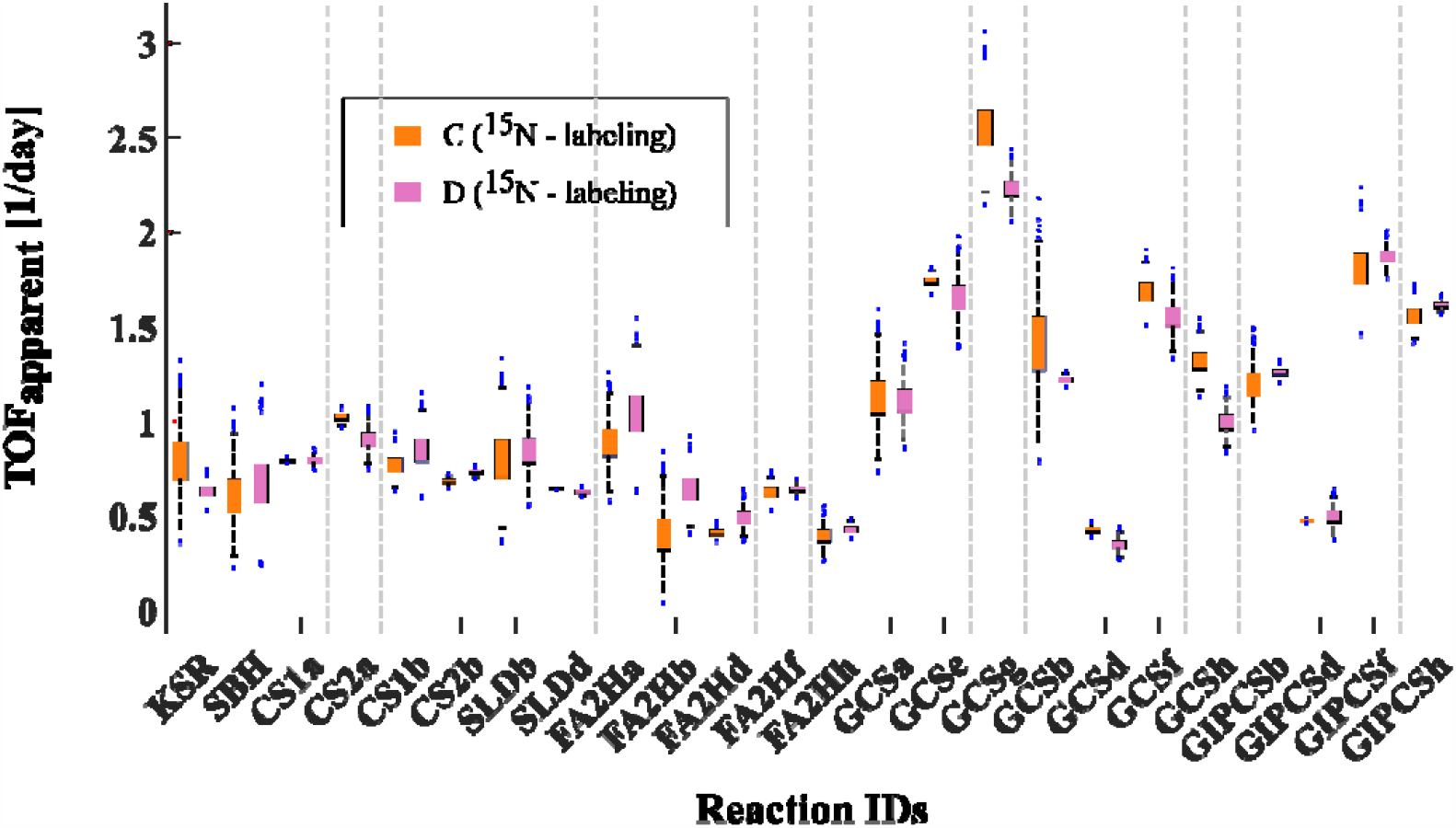
Apparent Turnover frequency (TOF) distribution across sphingolipid species. This figure illustrates the derived apparent turnover frequencies (TOF or values) for various sphingolipid species. Linear regression of the nonlinear model curve – was employed to determine these distributions. Rapid turnover is evident in GlcCer and GIPC species, indicating their pivotal role in dynamic cellular processes. In contrast, other species demonstrate relative stability. Distributions are determined for species in biological duplicates (representative cultures C and D).

### Sphingolipid metabolic turnover rates

The dynamics of *de novo* sphingolipid biosyntheses turnover were explored using the metabolic pathway, as outlined in our previous work (Alsiyabi et al., 2021) (see Fig. 2A and Supplementary Table 1). Utilizing the *in vitro* ^15^N-labeled quantification derived from HPLC/ESI-MS/MS analysis, we assessed the metabolic turnover during the experiment by comparing the abundance of ^15^N-labeled sphingolipids to the total pool. The resulting dynamic isotopologue fractions (or isotopic incorporation/enrichment) curves (e.g., species in Fig. 2C) demonstrate that labeled sphinganine (“d18:0”), associated with the restrictive uptake of serine, is integrated into sphingolipid before the exponential phase (specifically in the initial 24 hours). The sphingolipid then displays relatively consistent turnover rates. The employed Arabidopsis T87 cell culture, which is undifferentiated and photoautotrophic, possesses a short doubling time of about 2.7 days. Notably, barring a few exceptions, sphingolipids with dihydroxy LCB backbones exhibit isotopic incorporation curves that gradually stabilize, reaching a plateau around day 5 of the exponential phase.

We propose an indirect method suggesting that the *in vivo* turnover number of a metabolic reaction can be apparently determined by observing the rate and extent of labeled nitrogen synthesis or integration. This method offers a practical way to gauge *in vivo* catalytic speed and efficiency, especially when direct assessments are problematic. Leveraging the first-order kinetic growth/decay model and regression analyses, our intention is to identify relative turnover numbers of the species *de novo* synthesis. Although metabolic networks inherently possess complexities, our findings indicate minimal impact from factors like enzyme compartmentalization and substrate availability. This assertion is supported by our study of ^15^N abundance relative to the complete nitrogen pool, which demonstrated little complications concerning substrate availability, with the exception of d18:1-hC24. The observed ^15^N accumulation and isotopic incorporation hint at *de novo* synthesis for nearly all species. Our modeling approach assumes a first-order exponential correlation between labeled nitrogen incorporation ratio and turnover rates. Consistent with this premise and findings from isotopic incorporation (Fig. 2C and 3), an exponential rise in m + 1 isotopologue fractions was evident for most sphingolipids until the mid-exponential phase (Fig. 2C). However, Supplementary Fig. 3 suggests that the total active pools might not always follow an exponential trajectory, as seen with t18:1-hC24. A closer look indicates that for this exception, ^14^N is simply substituted with ^15^N, with the overall pool size remaining relatively constant (see Supplementary Fig. 2).

Variations observed in technical replicates of each biological sample necessitated a specific analysis approach. This method was chosen to accurately delineate the pronounced intra-sample variability inherent in our dataset. Notably, significant variability, as depicted in Fig. 2C’s incorporation ratio bands, led to the adoption of the bootstrapping technique for subsequent analyses. This technique quantifies within the standard deviation across replicate measurements, highlighting the dataset’s intrinsic variability. Further, turnover frequency distributions, represented as 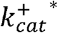 (or simply k), were derived using linear regression on the nonlinear model curve 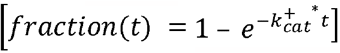 as shown in Fig. 3. It’s noteworthy that while glucosylceramide (GlcCer) and – glycosyl-inositolphosphoceramide (GIPC) species demonstrate rapid turnover, indicative of their role in dynamic cellular mechanisms, other species exhibit relative stability. Referring to *de novo* synthesis of the blueprint pathway, clusters of TOF 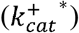 values emerged among particular sphingolipid pairs, notably between GCSf/h and GIPCSf/h. Both pairs metabolize t18:1-hC16/C24 to yield GlcCer t18:1-hC16/C24 and GIPC t18:1-hC16/C24, respectively. These 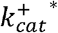 values indicate the heightened metabolic demands of the t18:1-hC16/C24 species, suggesting a pivotal role in maintaining cellular homeostasis of *de novo* synthesis.

Elevated accumulation levels of substrate d18:1-hC24 were noted, attributed to the activity of GCSg, which appears to function as an intrinsic shunt (shown in Supplementary Fig. 3). However, the confidence in incorporation data for this substrate is limited (see Fig. 2C), prompting caution in deriving universal conclusions about its regulatory efficacy. Furthermore, it is pertinent to mention that the cell-line used in this study lacks essential hormones for cellular differentiation. This observation is therefore consistent with prior research that postulates the critical role of GCS in mediating cell-type differentiation in Arabidopsis thaliana (Msanne et al., 2015). Further investigations are warranted to elucidate this finding.

Notably, the 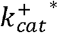 distribution associated with fatty acid hydroxylation reactions, symbolized by FA2H in Fig. 3, provides insights into the interactions within the mid-stream sphingolipid class, specifically the hydroxy-Ceramides (hCer). FA2Ha, which converts d18:0-C16 to its analogous hCer species, shows elevated activity. This activity could be linked to the increased concentrations of Cer d18:0-C16, a terminal species implicated in initiating programmed cell death. Elevated levels of Cer d18:0-C16 were observed in this culture (Fig. 2B and 3). A parallel phenotype was observed in an alternative experiment with the same cell line, reaffirming its inherent nature and negating the possibility of contamination. The LCB-Δ8-desaturase exhibited a preference for turnover towards the C16 fatty acyl over the C24 fatty acyl group (see SLDb in Fig. 3). Activities of Class I ceramide synthase (CSIa&b) were observed to be roughly equivalent, implying a competitive interplay between pathways. Specifically, CS1b’s pace seems modulated by the reaction that supplies it, SBH, which in turn influences its competition with CS1a and CSIIa. However, CSII displayed a marked preference for the C24 fatty acyl group, especially CSIIa. This suggests a regulatory role for CSIIa, particularly when SBH exhibits low turnover rates.

Additionally, consistency in 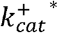 distributions was observed across biological replicates, with an average bhattacharyya’s coefficient (Bhattacharyya, 1943, 1946) overlap of 51.89+/-30.10%, across the reactions. We observed the apparent turnover frequency which is also a measure of the maximal catalytic efficiency 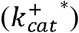 associated with LCB C4-hydrolase (SBH) and LCB Δ8 desaturase (SLDb) assumed to produce unsaturated LCBs bound to t18:0 ceramides and not free LCBs (M. Chen et al., 2012) had significantly high overlaps >94% between both biological replicates. This agreeably points to the essential role that SBH plays in branching di- and trihydroxy LCBs. Such overlap accentuates the credibility and rigor of these findings. Furthermore, the distribution of 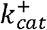 values associated with the activated CSII and GCS pathways also displayed significant differences between the two replicates which would suggest the modulatory role these paths play.

Moreover, the notable high metabolic demands for GIPC species, particularly those with trihydroxy LCBs, t18:0 and t18:1, reflected in the high values of GIPCSf and GIPCSh, underline the importance of GIPC and its prevalence in the lipid metabolism and membrane integrity of Arabidopsis (Liu et al., 2021). This aligns with the established and anticipated dominance of tri-hydroxylated sphingolipids in plants, specifically monounsaturated and saturated LCB derivatives (t18:1 and t18:0) (Buré et al., 2014; Liu et al., 2021). Interestingly, a consistent balance in the ratios of *de novo* pools for t18:1-hC24 to t18:0-hC24 for both hCer and GIPC was observed, approximately ∼1.5 fold, indicating the essential nature of these species and the delicate equilibrium needed for maintaining cellular homeostasis. This also suggests the crucial role of SLD, particularly SLDb and SLDc.

#### Dissecting dynamic shifts in metabolic flux through dynamic metabolic flux analysis (DMFA)

Analysis of reaction fluxes over time revealed key processes crucial for growth, proliferation, and cellular homeostasis. Our study employed an enhanced r-DMFA modeling, incorporating cubic spline interpolation and Hessian matrix regularization, to effectively dissect dynamic shifts in metabolic fluxes and bolster the robustness and flexibility of r-DMFA fitting (see Fig. 1 for workflow and see Materials and Methods). Transient fluxes were determined using ^15^N-labeled quantification from HPLC/ESI-MS/MS analysis as a relative measure of *de novo* sphingolipid biosynthesis. The metabolic pathway model adopted for this DMFA analysis aligns with the one described in our previous work (Alsiyabi et al., 2021) (see Fig. 2A and Supplementary Table 1). Similar to the referenced work, our model encompasses a thorough knowledgebase of all active metabolic transformations in Arabidopsis sphingolipid biosynthesis, accounting for enzyme promiscuity, parsimonious principle of minimal enzyme activity in most tissues, and the dominance of either C16 or C24 fatty acids in sphingolipids (Markham et al., 2006, 2011). We also adopted three specific assumptions from the referenced study, including i) LCB C4 hydroxylase (*sbh* gene) primarily hydroxylates free long chain bases, ii) fatty acid α-hydroxylase (fa2h gene) targets only ceramide-bound fatty acids, and iii) LCB Δ8 desaturase (*sld* gene) produces unsaturated LCBs bound to ceramides. By adopting these assumptions based on previous research, we minimize the degrees of freedom. Additionally, akin to the referenced study, our model consolidates certain metabolic pathway steps due to a lack of relative kinetics information and excludes reactions converting non-hydroxylated ceramides into complex forms, consistent with the composition of most glycosylated sphingolipids.

Utilizing the r-DMFA framework, transient *de novo* fluxes of internal reactions in the network were predicted and are illustrated in Fig. 4. These values were normalized to Serine Palmitoyl Transferase (SPT) to demonstrate how the flux through this reaction branches into different sphingolipid products. Metabolite pool sizes were also determined as an output of the DMFA fitting procedure (see Materials and Methods). Interestingly, the majority of the produced Long Chain Bases (LCBs), approximately 93%, are directed through class II ceramide synthase to generate FA containing ceramides, particularly those with more of C24 (VLCFA) and t18:0-LCB. Furthermore, the predictions indicate that the newly synthesized species undergo recycling only after reaching the mid-exponential phase, as evidenced by the negative fluxes observed subsequently. The decision not to recycle these newly synthesized species until the mid-exponential phase suggests a strategic metabolic shift that may be essential for cellular functions, potentially signaling a switch from biosynthesis to recycling that aligns with cellular demands for homeostasis and regulatory compliance. This could have profound implications for our understanding of sphingolipid dynamics in health and disease, where such regulatory mechanisms may be pivotal.

**Fig. 4.**
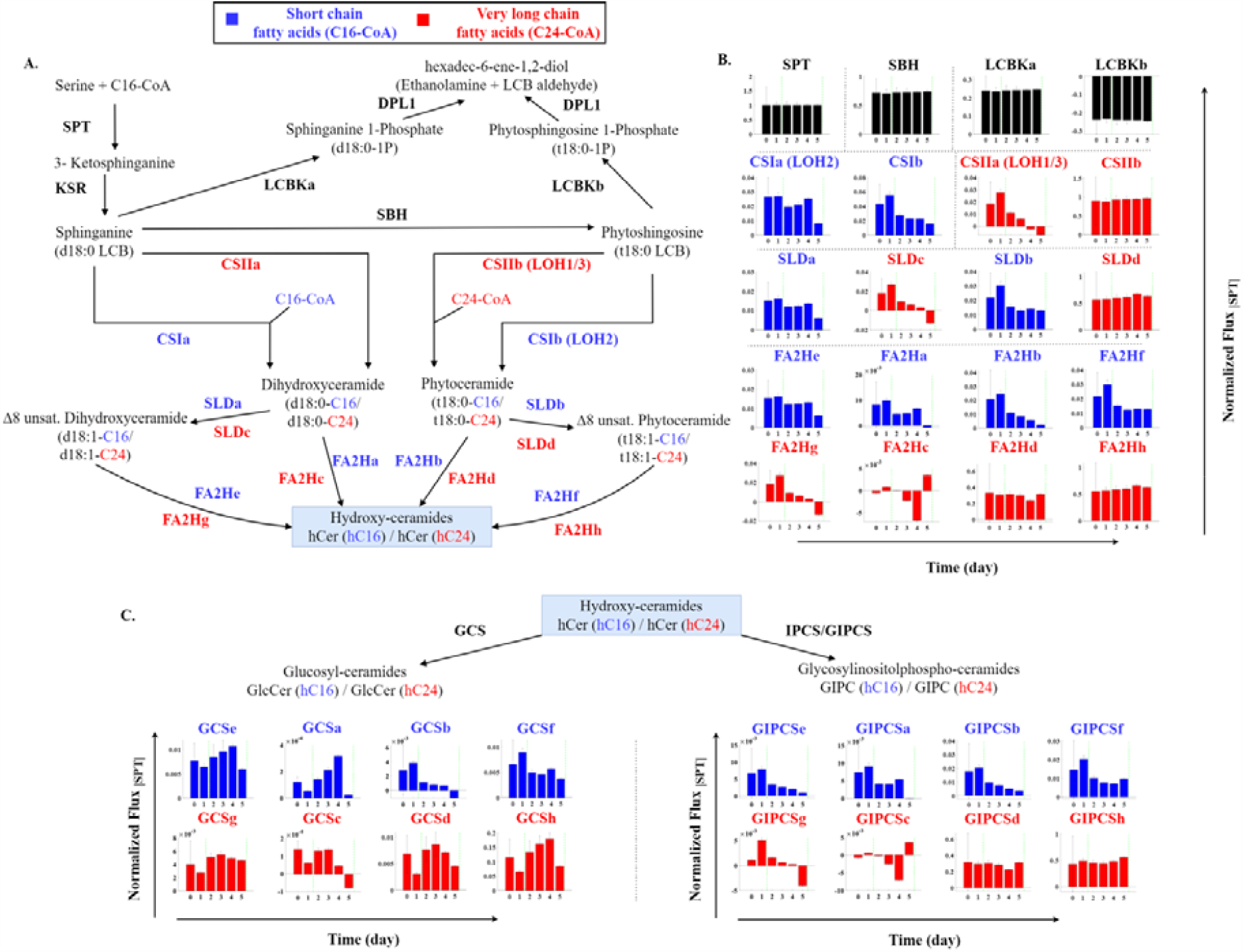
Transient *de novo* fluxes within the sphingolipid pathway network. **(A)** A metabolic network map used for r-DMFA analysis, and upstream of the hydroxy-ceramides (hCer) metabolite junction. This highlights internal reactions of the sphingolipid pathway **(B)** Corresponding transient fluxes from the pathway shown in (A). **(C)** Transient fluxes for processes occurring after the hCer. Overall, the flux values, normalized to serine palmitoyltransferase (SPT) activity, depict the branching of flux into different sphingolipid products. A predominant channeling of long chain bases (LCBs) (∼93%) through class II ceramide synthase for the production of very long-chain fatty acid (VLCFA) containing ceramides, particularly with C24 and t18:0-LCB, is observed.

To delve deeper into the enzyme requirements needed by the cell to match the experimental observation and maintain the observed wild-type (WT) flux distribution, the enzyme demand was calculated as a function of metabolite levels and transient *de novo* synthesis fluxes, *v* (Noor et al., 2016). This calculation provided a valuable measure of understanding resource allocation. The relative enzyme cost distribution computed is depicted in Fig. 5A. This analysis clearly revealed a significantly low level of CS1 (LOH2), and GCS enzyme activity, suggesting a close regulation of the enzyme’s function in response to substrate availability and metabolic demands, which results in reduced specific activity. However, it is plausible that other enzymes in the pathway may compensate for this deficiency, ensuring the overall efficiency of the biochemical process (Noor et al., 2013). Furthermore, this relatively low enzyme activity warrants additional investigation to determine if environmental stressors or nutrient limitations could instigate higher demands, thereby linking these enzyme activities to an adaptive response aimed at conserving energy and resources.

**Fig. 5.**
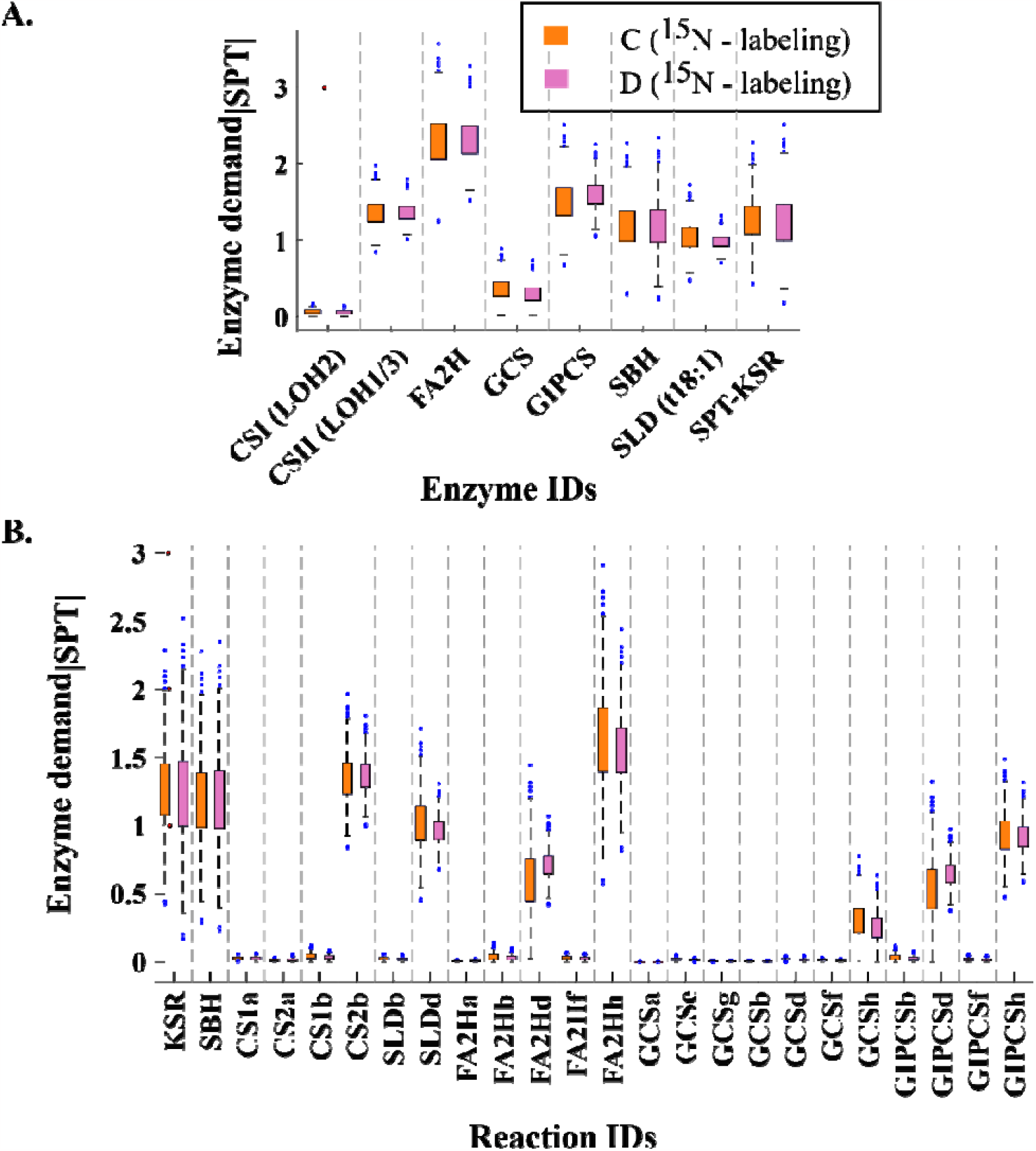
Analysis of enzyme demand and relative cost distribution in the sphingolipid biosynthesis pathway. **(A)** Distribution of relative enzyme costs, showcasing the energy and resource allocation within the metabolic network normalized to the SPT limiting enzyme. The analysis reveals significantly low activity levels of CS1 (LOH2), and GCS enzymes, indicating tight regulation in response to substrate availability and metabolic demands. The relative enzyme activity levels, in relation to the first and rate-limiting step catalyzed by SPT, suggest key roles for specific enzymes in maintaining pathway homeostasis and reaction rates. **(B)** Enzyme specificity investigation, representing the demand of each enzyme concerning each reactive sphingolipid species, highlighting the interplay of enzymes with various lipid species, and revealing potential regulatory hotspots. The figure highlights the variability in enzyme cost specificity, with notably high demands for FA2Hd and FA2Hh, enzymes involved in the conversion of t18:0-C16 and t18:1-C24 to their respective analogous hydroxyceramides (hCers), underscoring the pivotal role of FA2H in meeting the metabolic demands of t18:0/t18:1-hC24 species.

As depicted in Fig. 5A, the enzyme cost for most of the biochemical transformations associated with biosynthesis (*de novo*) is relatively at par with the first and rate-limiting step of the pathway, the reaction catalyzed by SPT (Luttgeharm et al., 2016). This observation underscores the essential roles of SPT, SBH, and CSII (LOH1/3) in the thermodynamic feasibility of *de novo* synthesis. It also suggests that LCB branching at SBH may play a pivotal role in maintaining homeostasis within the sphingolipid pathway (Noor et al., 2013). Moreover, this bottleneck in enzyme demands, which can directly infer thermodynamic bottlenecks, indicates these enzymes are key players required to maintain the measured reaction rates and/or cellular homeostasis.

Nonetheless, an almost twofold relative cost was observed for FA2H, which is connected to the low turnover numbers observed in Fig. 3. This wide range of distribution for FA2H also suggests that reactions involving this enzyme have a broad range of positive substrate/product concentration ratios, which can result in an overall forward flux of the reaction (Noor et al., 2013). This implies that a closely regulated mid-stream enzyme like FA2H may be a key player in *de novo* synthesis. Subsequently, a two-sample Kolmogorov-Smirnov test (KS test) (Massey, 1951) was performed to determine if there were any statistically significant differences in the distributions of the calculated enzyme costs computed for experimental biological replicates. Interestingly, a significant difference (p < 0.001) was found for four enzyme classes, namely CS1 (LOH2), GCS, GIPCS, and SLD (t18:1). Visual inspection of these distributions (data not included but can be inferred from Fig. 5) revealed that one replicate is a subset of the other, with calculated average Bhattacharyya coefficient overlap of 95.90 +/-3.91%.

Further examination of enzyme specificity was conducted by computing each enzyme demand concerning each reactive sphingolipid species, as shown in Fig. 5B. This analysis revealed enzyme cost specificity with the highest visible variability in the FA2H class of enzymes. Notably high demands were observed with FA2Hd and FA2Hh, which convert t18:0-C24 and t18:1-C24 to their respective analogous hCers. This observation underscores the intrinsic role of FA2H in meeting the heightened metabolic demands of the t18:0/t18:1-hC24 species. Additionally, in general, enzyme specificities towards C24 VLCFA were observed to be higher than those towards FA C16. The average Bhattacharyya coefficient overlap for the biological replicates across the reactive species was 90.21 +/-7.80 %. Interestingly, the low demand of GCS enzyme agrees with previous studies’ postulates linking GCS to cellular differentiation which our cell-line lacks (Msanne et al., 2015).

Building on these observations, a uniform convex polytope sampling strategy was employed to thoroughly analyze the high-dimensional dynamic flux solution spaces, considering different alternate optimal dynamic flux distributions. This involved uniformly drawing representative random transient concentration profiles within the given experimental constraints to truncate the bounded flux polytope, a method commonly used for characterizing the solution spaces of metabolism (Amaran & Sahinidis, 2012; Herrmann et al., 2019; Schellenberger & Palsson, 2009) (see Materials and Methods). Further, we examined a hypothesis, involving the partitioning of the sampled dynamic flux distribution set within the metabolic network. This hypothesis suggests that the relative flux modulation ratios, defined as the fold changes of each flux measurement relative to the mean flux of the distribution, directly influence the biological partitioning of reactions. Specifically, it was hypothesized that reactions with high variations in the distribution of flux modulation ratios would display a distinct partitioning pattern compared to those with low variations in the distribution of modulation ratios, thereby indicating a potential regulatory mechanism in the distribution of metabolic flux.

Moreso, cellular homeostasis, which is essential for the proper functioning of cells, is linked to the tight regulation of metabolic pathways. Fig. 6A and 6B provide insights into how variations in flux modulation ratios across different biological replicates could be linked to the maintenance of cellular homeostasis. Specifically, the red cluster/partition, characterized by low variations in flux modulation ratios, encompasses reactions that are tightly regulated and likely constitute the limiting set of the biosynthesis (*de novo*). The identified candidate reactions (CSIIb, SLDd, FA2Hh, and GIPCSh) are in line with the activities of known essential enzymes such as SPT, KSR, and SBH. This tight regulation is crucial for maintaining cellular homeostasis, as it ensures that the synthesis of key cellular components is carried out efficiently and consistently, even under varying environmental conditions. On the other hand, the green and blue clusters/partitions, characterized by moderate to high variations in flux modulation ratios, consist of metabolic invariant sets that have little to no tight regulations on *de novo* synthesis (see Fig. 6C and 6D boxplot distributions). These reactions, while not as tightly regulated, still play a vital role in cellular homeostasis by providing flexibility and adaptability to the metabolic network, allowing cells to respond to changes in the environment while maintaining their internal equilibrium. Together, these findings highlight the critical balance between tight regulation and flexibility in the metabolic network, which is essential for maintaining cellular homeostasis and ensuring the proper functioning of cells.

**Fig. 6.**
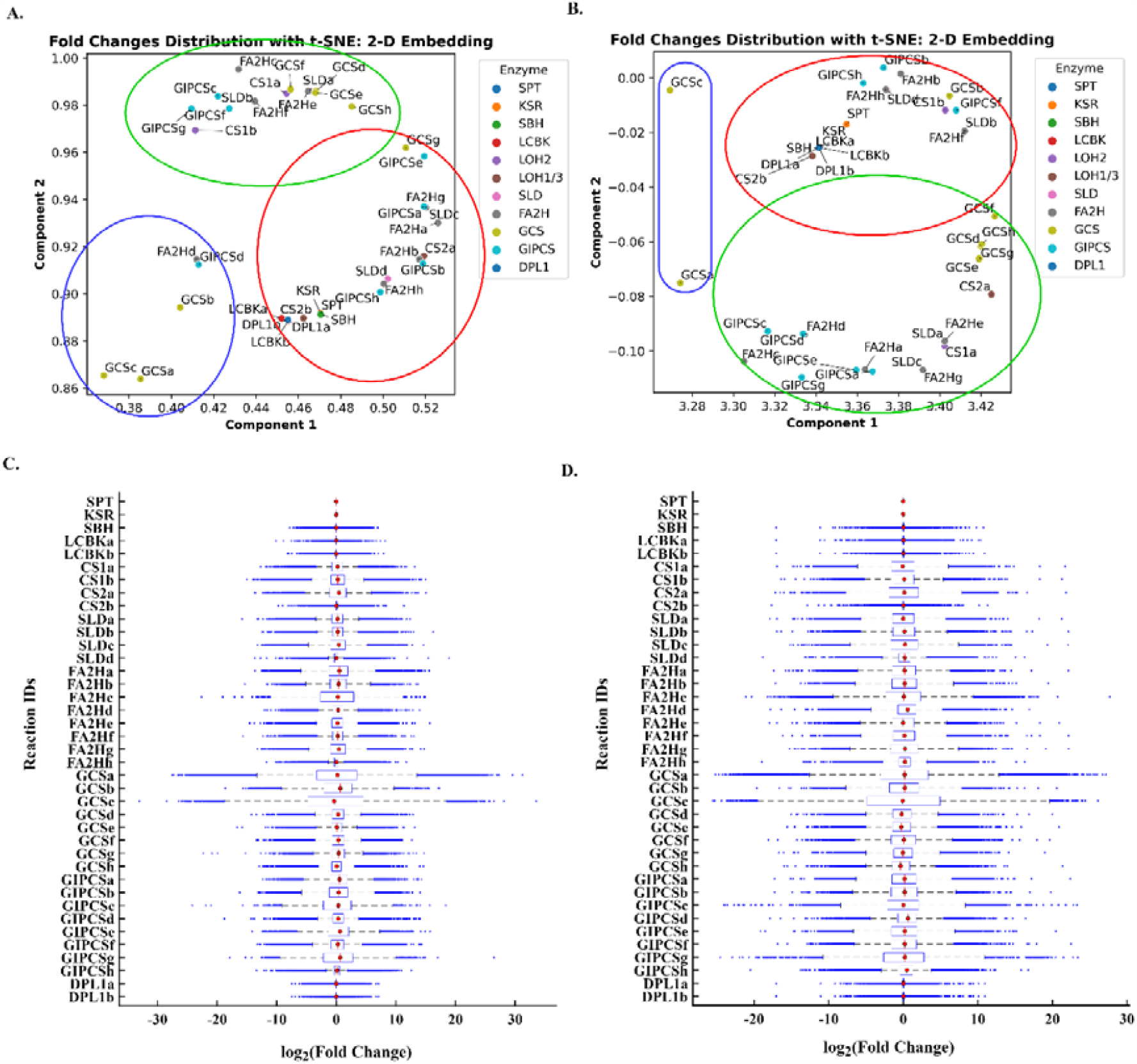
Insights into metabolic pathway regulation through flux modulation ratio analysis. **(A) & (C)** Scatter plot and boxplot distribution for the first biological replicate, respectively. These depict variations in flux modulation ratios, highlighting tightly regulated (red cluster) and flexible metabolic reactions (green and blue clusters). The figures represent three distinct partitioning of flux modulations. **(B) & (D)** Scatter plot and boxplot distribution for the second biological replicate. The boxplot distributions of flux modulation provide a view of metabolic congruence or divergence. While there are slight differences between the two replicates, these are deemed insignificant and are likely due to noise effects in the measurements. Key reaction players remain consistent across both replicates.

Additionally, a customized MATLAB implementation of the r-DMFA modeling framework was developed to precisely predict transient metabolic reaction fluxes following enzymatic perturbations as knockout (KO) (see Materials and Methods). This *in silico* KO/essentiality hypothesis testing approach could potentially minimize extensive experimental iterations in subsequent investigations and validations. This process was repeated for each enzyme class and reaction specificity and can be extended to probable enzymatic perturbations. As briefly depicted in Fig. 1B, the experimentally fitted and transient metabolomics result in the predicted transient reaction fluxes for the reference (wild type). The obtained WT profiles are then used to generate dynamic models whose metabolic responses/profiles are screened with respect to different perturbations.

#### *In silico* metabolic analysis of single enzyme deletions and reaction knockouts perturbations

To evaluate the metabolic network’s functionality without any additional regulatory interactions, we induced perturbations via single enzyme and reaction knockouts. We then leveraged the constrained based r-DMFA framework, in similitude of established global minimization of metabolic adjustment (MOMA) principle (Segrè et al., 2002), to predict each dynamic model’s response to the induced perturbation. As an objective here, we compared the predicted responses from single knockouts with experimental wild-type predictions, such that the r-DMFA framework, utilizing quadratic programming and relaxing optimal wild-type flux assumptions, more accurately predicts metabolic phenotypes of mutants and assists in comprehending cellular adaptation to enzymatic dysfunction. This approach leverages on the convex transformation methods employed in the global optimization algorithm of BARON solver (Nohra & Sahinidis, 2018). The findings detailed here originate from *in silico* simulations, critical for essentiality predictions. Before initiating laboratory experiments, it was vital to ensure that our methodology could foresee the outcome of anticipated experiments (Schellenberger et al., 2012). Our approach’s credibility was preliminarily confirmed by aligning our *in-silico* essentiality predictions with prior studies and corroborating the functions of the pinpointed enzymes/genes with literature-documented experiments. Subsequent research may necessitate further validation via laboratory experiments.

The metabolic network model was first examined through the lens of single enzyme deletions or knockout (KO) perturbations. By incorporating specific constraints into the model and applying an evaluation metric, *z*_*d*_, normalized to the maximum distance on a logarithmic scale (as depicted in or knockout (KO) perturbations. By incorporating specific constraints into the model and applying an Fig. 7A and detailed in the materials and methods), the metabolic distance between each *in-silico* mutant dynamic flux phenotype and the WT dynamic flux phenotype was assessed using Spearman’s rank correlation, as shown in Fig. 7B. This approach was essential to understand the role of each enzyme within the metabolic network and to pinpoint those critical for the wild-type phenotype. The analysis in Fig. 7A highlighted the significant roles of enzymes such as SPT, KSR, LOH1/3 (or CS II), FA2H, SLD, and GIPCS, which displayed pronounced metabolic distances from the WT; with z_*d*_ values greater than 0.5-fold of the limiting SPT enzyme activity. LOH2 (or CS I) and GCS had moderate z_*d*_ values around 0.5-fold.

**Fig. 7.**
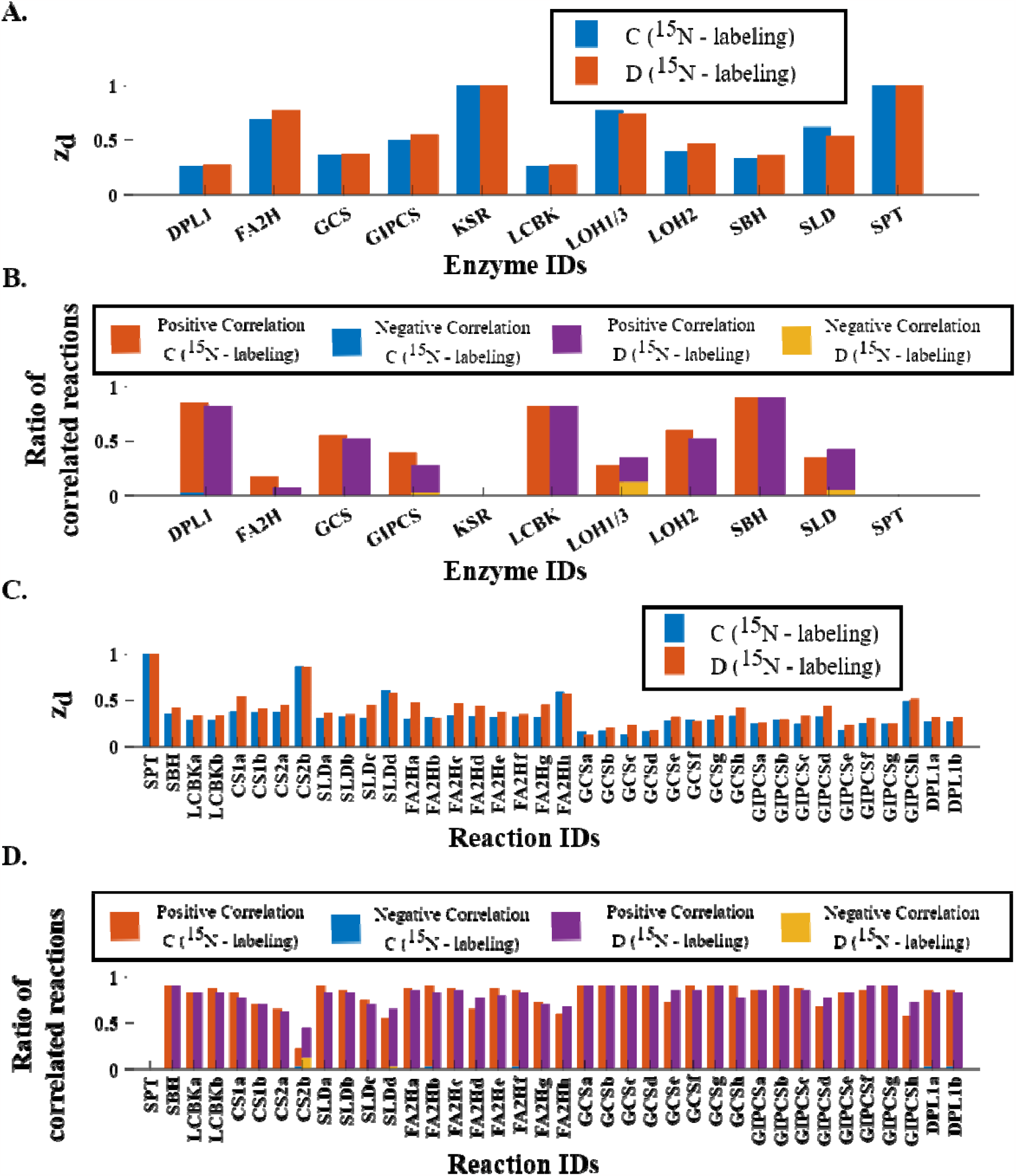
*In silico* Metabolic Analysis of Single Enzyme Deletions and Reaction Knockouts. **(A)** Evaluation of the metabolic network model through the lens of an evaluation metric,, normalized to the maximum distance on a logarithmic scale. The approach using single enzyme deletions or knockout (KO) perturbations was undertaken to discern the role of each enzyme within the metabolic network. Here, key enzymes such as SPT, KSR, LOH1/3 (or CS II), FA2H, SLD, and GIPCS are highlighted for their pronounced metabolic distances from the WT phenotype, with special note on the moderate values exhibited by LOH2 (or CS I) and GCS. **(B)** Spearman’s rank correlation analysis, assessing the metabolic distance between each *in silico* mutant dynamic flux phenotype and the WT dynamic flux phenotype. The panel further visualizes the ratios of reactions that show both positive and negative correlations between the in-silico mutant and WT profiles, yielding insights into metabolic alignment or divergence. Enzymes with pronounced metabolic divergence are highlighted, and the potential reversible action of the long-chain-base kinase (LCBK) and its modulation of SBH function is discussed. **(C) & (D)** extension of the analysis by focusing on single reaction knockouts. Panel C delves into the metabolic distance analysis, while Panel D concentrates on correlation analysis at the reaction level. These panels aim to pinpoint crucial reactions within the metabolic network, emphasizing those that, upon perturbation, induce significant deviations from the WT dynamic flux phenotype. Observations such as the reconfirmation of enzyme specificities and the interconnectedness of enzyme abundance with specificity in relation to reactive species in the metabolic network are presented. Also, the specificity towards C24 VLCFA over C16 FA and the increased concentration of ceramide d18:1-hC24, implying a reduction in LCBs due to a diminished rate through SPT, are particularly emphasized.

Interestingly, the SBH mutant exhibited a minimal (<0.5-fold) metabolic distance to the WT, contradicting its traditionally understood role in cellular stability. Its deficiency is projected to lead to metabolic distant phenotypes and an increased potential for programmed cell death (PCD) due to sphinganine (d18:0) accumulation (M. Chen et al., 2008). This observation necessitated further examination of flux activities. Additionally, ratios of reactions showcasing both positive and negative correlations between the *in-silico* mutant and WT profiles were calculated, as illustrated in Fig. 7B. This data provided insights into metabolic congruence or divergence. The observations confirmed the prominent roles of the aforementioned limiting mutants, with significantly low ratios (<0.5). However, a stimulated metabolic divergence was observed for CS II or LOH1/3, GIPCS, and SLD mutants, evident in the relative increase of negatively correlated reactions when compared to the WT profiles. Moreso, the low ratios observed in Fig. 7B for FA2H, GCS, CS I and CS II as well as SLD mutants suggest significant metabolic reprogramming indicative of synergistic role these enzymes may play along this pathway.

Fig. 7B also suggests the buffering activities of LCBK, DPL1, and SBH mutants having high ratio of significantly positive correlated reactions at par with the observations from their z_d_ values in Fig. 7B. Given the observed low ratio of negatively correlated reactions in the DPL1 mutant, it suggests that DPL1 might be involved in some metabolic reprogramming Upon further scrutiny, a reversible action of the long-chain-base kinase (LCBK), potentially a putative sphingoid-phosphate phosphatase (LCBPP1), emerged as a key modulator of SBH function (data not shown here). The long-chain-base kinase (LCBK) catalyzes the phosphorylation of long-chain bases to form bioactive long-chain base phosphates. These phosphorylated species modulate cellular growth and differentiation. On the other hand, the putative sphingoid-phosphate phosphatase (LCBPP1) is believed to dephosphorylate these molecules, restoring them to their long-chain base precursors. Additionally, it was identified as a potential rate-limiting enzyme in the degradation of phosphorylated phytosphinganine (LCB-1Ps) via the competing path of dihydrosphinganine-1-phosphate lyase (DPL1) (Lambour et al., 2022; Nishikawa et al., 2008; Tsegaye et al., 2007). This observation hints at a regulatory interaction that could either serve as a recycle shunt or play a limiting role in degradation. The differential activity of LCBK underscores its importance in modulating SBH function and in the degradation of LCB-1Ps. This regulatory interaction could be vital for *de novo* synthesis and dephosphorylation of LCB-1Ps by phosphatases (LCBPP1/SPP1). However, additional experimental validation is necessary to decode the reversible roles of these putative enzymes and their implications on *de novo* biosynthesis.

Two earlier studies (Imai & Nishiura, 2005; Worrall et al., 2008) have explored this interaction inconclusively, suggesting a potential coordinated regulation between LCBK and its reversible counterpart. This underscores the need for more comprehensive research to thoroughly understand the interactions between these enzymes and their collective role in maintaining cellular homeostasis. LCB-1-phosphates (LCB-1Ps) are recognized as lipid messenger molecules implicated in plant stress signaling (Guo & Wang, 2012). The most recent investigation by (Nakagawa et al., 2012) explored the functional role of LCBP phosphatase in regulating LCBP levels and its participation in the stomatal response to abscisic acid (ABA) signaling pathways in Arabidopsis, with potential implications for enhancing water use efficiency and drought resistance. This further highlights the dynamic balance of LCBs/LCB-1Ps. Moreover, the enzymes LOH (2, 1/3), FA2H, and GCS were found to have a moderate impact on the metabolic network, with their influence increasing in the order mentioned. Notably, GCS, despite having a low enzyme cost in the WT, exhibited a significant impact on the metabolic network. This observation is intriguing as it implies that even enzymes with lower costs in the wild-type phenotype can have a substantial impact on the metabolic network, underscoring the complexity and interconnectivity of the system. This highlights the need for a comprehensive analysis that considers not only the individual enzyme costs but also the overall impact of each enzyme on the metabolic network. Understanding the relative importance of these enzymes could provide valuable insights into potential targets for metabolic engineering.

The analysis of enzyme specificity regulations was extended to single reaction knockouts, as depicted in Fig. 7C and 7D, which highlight both the metabolic distance and correlation analysis at the reaction level. This layer of analysis aimed to pinpoint critical reactions within the metabolic network that, when perturbed, induce significant deviations from the wild-type dynamic flux phenotype, akin to the analysis at the enzyme level. Trends analogous to those observed in the enzyme level knockout were uncovered. A notable observation was the reconfirmation of enzyme specificities; the specificities towards C24 VLCFA were found to be higher than those towards FA carbon length of 16, C16. Moreover, the activity of SLDc bolstered the experimental observation that the concentration of VLCFA-containing ceramide d18:1-hC24 increases, thereby suggesting a reduction in the overall amount of LCBs produced due to the diminished rate through SPT. This reiteration underscores the interconnectedness of the enzyme abundance and specificity with respect to reactive species in a metabolic network. Understanding the regulations behind this balance will be key to engineer crops with improved yield while enabling robust response to biotic and abiotic stress. Reinforcing our observations on enzyme abundance and specificity, we re-hypothesize that enzyme specificity plays a pivotal role in the rapid evolution and regulation of *de novo* synthesis, as evidenced by dynamic flux distribution in the metabolic network, guiding crucial metabolic regulatory shifts.

With the advancements in high-throughput metabolomics and *in silico* modeling, dynamic models of metabolism can enhance our understanding of the metabolic regulation of the sphingolipid biosynthesis pathway. The r-DMFA framework can be applied to other systems to study the combined regulatory impacts of biosynthesis and catabolism of metabolic reactive species over time. In future, this dynamic modeling framework will produce more experimentally testable hypotheses regarding regulation in sphingolipid biosynthesis pathway and thereby minimize exhaustive experimental effort.

## Conclusion

The dynamic metabolic trajectory of enzymes and reactions within the sphingolipid biosynthetic pathway in Arabidopsis offers a captivating look into cellular metabolism. Through a series of meticulous *in silico* analyses, regulatory mechanisms underpinning this pathway were elucidated. The metabolic turnover rates established the pivotal roles of several enzymes, including SPT, KSR, LOH1/3, and FA2H. Furthermore, dynamic metabolic flux analysis (r-DMFA) showcased the branching nature of the metabolic flux, emphasizing the significant roles of particular enzymes in the wild-type phenotype. GCS enzyme was identified to be tightly regulated with low enzyme costs. Meanwhile, *in silico* metabolic analysis of enzyme deletions and knockouts provided a closer look at the network’s functionality. With this, the dynamic flux estimates suggested that long-chain-base kinase (LCBK) mediated sphingolipid-base hydroxylase (SBH) activity shortage and may be a potential regulatory scheme. Additionally, the role of SBH, LCBK, FA2H, CS II, and GCS, in ensuring cellular viability and managing programmed cell death, is meticulously highlighted, opening avenues for further research in the biosynthesis and catabolism of sphingolipids. Notably, enzyme specificity towards the phytosphingosine-based (tri-hydroxylated) groups and VLCFA emerged as a critical component, influencing the overall metabolic trajectory. With the integration of high-throughput metabolomics and advanced *in silico* modeling, such analyses present a promising avenue for generating experimentally testable hypotheses. This research not only illuminates the inner workings of the sphingolipid biosynthesis (*de novo*) but also sets the stage for future endeavors in metabolic engineering, potentially improving crop yield and stress response mechanisms.

## Materials and methods

### Plant material and culture conditions

This study utilized undifferentiated and photoautotrophic *Arabidopsis thaliana* T87 cells originated from the Columbia ecotype, Col-0. Cells were cultured in NT-1 liquid medium (Murashige and Skoog medium enriched with vitamins, 30 g/l sucrose, 1 mM KH_2_PO_4_, 1 mg/l thiamine, 100 mg/l myo-inositol, and 2 μM 2,4-dichlorophenoxyacetic acid; pH 5.8 adjusted with KOH) in 100 ml flasks. Cultures were maintained at 22°C, 55% humidity, agitated at 120 rpm, and continuously illuminated at a photosynthetic photon flux density (PPFD) of 100 μmol m^−2^ s^−1^.

For internal standard and dynamic labeling, T87 cells were cultured in ^15^N-enriched NT-1 liquid medium containing K^15^NO_3_ and ^15^NH_4_^15^NO_3_ (Cambridge Isotope Labs), under identical culture conditions. Standard sub-culturing was performed every 7 days, with 1 ml of the existing cell suspension transferred to 50 ml of fresh NT-1 medium. The ^15^N medium was introduced dynamically with daily sample collections initiating immediately and continuing for 6 days, up to the experimentally determined mid-exponential growth phase. A control culture was concurrently maintained in NT-1 liquid medium with the natural ^14^N isotope, under identical conditions. Each culture setup entailed two technical replicates with two biological replicates per setup.

To ascertain growth phases, growth rates were ascertained by freeze-drying and dry-weighing equal volumes of suspension cultures (bi-replicates) over a 15-day period. Data were then plotted on a semi-log graph, with linear regression applied to determine the growth rate from the slope of the best-fit line (see supplementary Fig. 1 for growth curve).

### Sphingolipids extraction and quantification

The method employed for sphingolipid extraction and quantification was adapted from Markham and Jaworski (2007) (Markham & Jaworski, 2007). The profiling methods which have been modified was published in Cahoon et al. (2021) (Cahoon et al., 2021). Specifically, 10-30 mg of lyophilized cells were homogenized and extracted employing a solvent mixture of isopropanol:heptane:water (55:20:25, v/v/v). Following extraction, the supernatants were dried and de-esterified using methylamine in a ethanol:water solution (70:30, v/v). The lipid extracts were then re-suspended in a solvent composition of tetrahydrofuran:methanol:water (5:2:5, v/v/v) with 0.1% (v/v) formic acid. Sphingolipid species analysis was conducted using a Shimadzu Prominence ultra-performance liquid chromatography system and a 4000 QTRAP mass spectrometer (AB SCIEX). Internal standards were incorporated for precise sphingolipid class identification.

For precise quantification of ^14^N / ^15^N labeled sphingolipids, standard curves were generated through sphingolipid (analyte) isolation and analysis utilizing the 4000 QTRAP® LC-MS/MS system across multiple dilutions. Specifically, the HPLC/electrospray ionization tandem mass spectrometry (HPLC/ESI-MS/MS) method, as described by Markham and Jaworski (Markham & Jaworski, 2007), was employed, predicated on the principles of ion transitioning, separation, and fragmentation. In this method, chromatographic separation was achieved using a reversed-phase C18 column, facilitated by a specified gradient elution of aqueous and organic mobile phases. Following separation, the eluent was subjected to electrospray ionization (ESI) to generate charged molecular ions, which were subsequently directed into the tandem mass spectrometer (MS/MS). Within the MS/MS, ions were initially filtered by the first quadrupole (Q1) based on their mass-to-charge ratios (m/z), then fragmented in the collision cell (Q2) to yield a spectrum of product ions, which were filtered and detected by the second quadrupole (Q3). The methodology hinged on the selection of optimal precursor-product ion transitions, accompanied by the meticulous optimization of mass spectrometer parameters including de-clustering potential (DP), collision energy (CE), and retention time.

Utilizing the HPLC/ESI-MS/MS technique, 168 distinct sphingolipid species/analytes (LCBs, ceramides, GlcCers, and GIPCs) werequantified over a designated time course. To dynamically trace ^15^N incorporation into sphingolipid species, analytes were scrutinized for base m/z (m + 0) and +1 m/z shift (m + 1) isotopologues. The (m + 0) isotopologue, e.g., 302.3 m/z for “d18:0”, denotes the mass wherein all atoms exist in their predominant isotopic form, predominantly ^14^N. In contrast, the (m + 1) isotopologue, e.g., 303.3 m/z for “d18:0”, represents the mass shift due to a single ^15^N integration.. The contribution of naturally occurring 13C isotopes is low --- only 1 out of 100 naturally occurring carbon molecules is 13C. The long-chain bases, ceramides and glycosylceramides measured here have fewer than 100 carbon atoms. Larger sphingolipids such as GIPCs are where the 13C isotope effect does create a small proportion of m+1 molecules that are heavy because of naturally occurring 13C. The incorporation of 15N is pronounced and easily distinguished from possible 13C isotope effects, especially in the smaller sphingolipid classes that contain only 18 to 48 carbons. Additionally, technical duplicates were executed for each biological replicate inclusive of cultures grown on ^15^N and control cultures on ^14^N. Data analysis and quantification were performed using Analyst 1.5 and MultiQuant 2.1 software, with considerations for exact mass and retention time as described in (Markham & Jaworski, 2007).

### Dynamic modeling

Dynamic metabolic modeling captures both the metabolic and regulatory state of a pathway through temporal reaction flux changes. Unlike many genome-scale metabolic models (GSMMs) which are static, dynamic metabolic models (DMMs) capture the dynamic nature of metabolic networks which is crucial for understanding and engineering biological systems. In this section, we highlight the *in silico* dynamic modeling approaches used to analyze the metabolic data and generate insights.

#### Estimating dynamic flux using regularized DMFA

A regularized form of the dynamic metabolic flux analysis (L-DMFA) framework by (Leighty & Antoniewicz, 2011) was used to predict the dynamic metabolic fluxes, *v*(*t*). DMFA methodology uses a DOA approach to fit concentration measurements directly and relies on non-steady state mass balances for metabolite pools. The model non-steady state mass balances for metabolite pools are represented by:

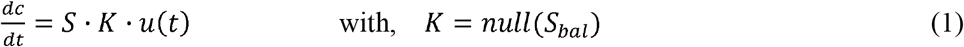

In this context and for a given metabolic network, we denote, as the stoichiometric matrix, *K* as the null space matrix of *S*, and *c*(*t*)as the vector of metabolite concentrations. Metabolites are categorized as balanced, typically intracellular, or non-balanced, typically external. Leighty and Antoniewicz, (2011) defined and determined internal free fluxes, *u*(*t*)as a function of linear splines. It divides the time domain [*t*_1_,*t*_n_]of a culture into smaller intervals [*t*_*k*−1_,*t*_*k*_], assuming constant rate changes in fluxes between distinct DMFA time points. Hence, the flux dynamics are modeled as linear flux changes between two points as:

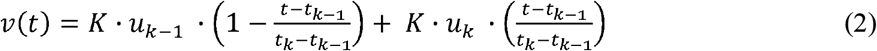

In a more compact form:

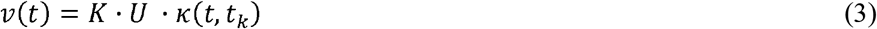

where, *U=*[*u* _*k*−1_,*u*_k_ *u*_*n*_ and *u*_k_ denote the free internal fluxes to be fitted at times *t*_*k*−1_ and *t*_*k*_ respectively. *k(t,t*_*k*_) is a time dependent parameter matrix for the linear spline with its collocations defined in (Leighty & Antoniewicz, 2011). Integrating equations (1) and (3) yields:

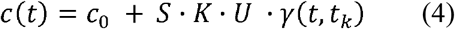

where, c_0_ is a fitted variable defined as the initial metabolite concentrations, and γ(*t,t*_*k*_) is the integral form of *k(t,t*_*k*_). For estimating flux transients, the variance-weighted sum of squared residuals (SSR) is minimized subject to equation (4) as follows:

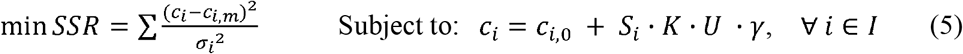

where, c_*i,m*_ are measured concentration profiles for metabolite pools; *I* are sets of metabolites in the model; *σ*_i_ are the specific measurement variances assumed as experimentally determined fluctuations. Hence, direct linear explicit and global optimums, *p* are obtained from:

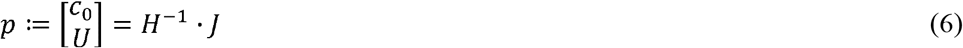

where, the Hessian and Jacobian matrices,*H* and respectively are calculated as:

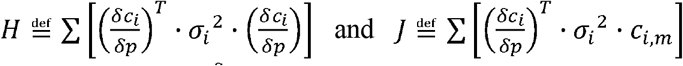

with the derivatives of 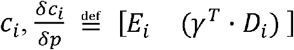
where, *D*_*i*_ = *diag*(*S*_*i*_·*K*) and 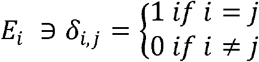

With the current formulation, solving the linear system involves inverting the Hessian matrix, *H* to estimating the parameter set,*p*. However, when data includes noisy (such as in our study the large variation in the distributions of σ_i_) or incomplete measurements the problem might be ill-posed; that is, it may not have a unique solution, or small changes in the data might lead to drastically different solutions. *H* could as well be near singular or ill-conditioned, its inversion can lead to numerical instability. Overfitting is another concern, where the estimated model fits the noise in the data rather than the underlying process. Tikhonov regularization of *H* was used to address these issues by adding a regularization term that conditions *H*, making its inversion more stable. This regularization promotes numerical stability and generalization The regularized Hessian, *H* ^′^ is given by:

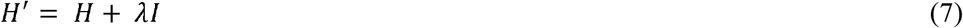

where, *I* is an identity matrix. With the regularized Hessian, *H* ^′^, the regularization parameter, λ, balances the trade-off between fitting the data and regularizing the solution in equation (6), and can be chosen via methods like cross-validation or information criteria. In this study, the regularization parameter, λ, is chosen through a systematic grid search optimization utilizing the SSR, and Bayesian Information Criterion (BIC). Optimal λ was chosen with the smallest BIC value.

Following the L-DMFA method that assumes linear flux changes between two DMFA time points, we use an iterative process to find the best set of linear time intervals that lead to a converging minimum of reduced SSR. However, we use cubic spline interpolation to include each data measurement point and generate additional points within these intervals. These additional points are treated as inflection points, which helps in assuming linear flux changes between them. This adjustment improves L-DMFA fitting by providing a smoother curve that accurately represents the non-linear changes in metabolic fluxes from the experimental data. Combined with the earlier regularization step, this approach strengthens the model’s robustness. Post this refinement, the formulation aligns with the L-DMFA approach as delineated by Leighty and Antoniewicz (2011), particularly in the technique employed to ascertain the standard deviations of the estimated metabolic fluxes.

The regularized DMFA framework, r-DMFA as a modified version of the MATLAB implementation developed by Leighty and Antoniewicz (2011) (Leighty & Antoniewicz, 2011) was scripted and used to calculate the reaction rates for each enzymatic reaction in the network.

#### Estimating enzyme metabolic demands

By intertwining thermodynamics and enzyme kinetics, we used the concept of enzyme metabolic demands which facilitates a holistic understanding of underlying mechanism dictating the efficiency, speed, and equilibrium of enzymatic reactions in our network. This proffers an estimation of active enzyme molecules partaking in the reaction. To this end, we determined the variable enzyme metabolic demands, *E*_*MD*_ for each reaction in the network as defined by (Noor et al., 2016). This was summed over all reactions to also obtain the enzyme level metabolic demand. For a single reaction, *j*, the enzyme metabolic demand is defined as the ratio of metabolic reaction flux, *v*_*j*_ (t) and the relative turnover frequency 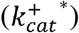, our experimentally fitted TOF and a measure of reaction speed, given by the equation:

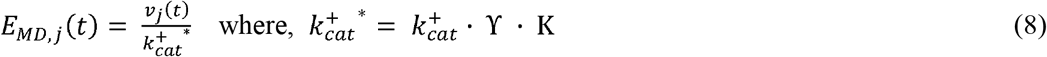

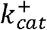 will be the actual enzyme turnover number independent of thermodynamic and substrate saturation. γ ∈[−1,1], is the thermodynamic driving force. *K* ∈[−1,1], is the enzyme specific substrate saturation factor.

#### Analyzing transient flux distribution through dynamic flux sampling

A concerted approach employing a uniform convex polytope sampling strategy was adopted to delve into the high-dimensional dynamic flux solution spaces, encompassing diverse alternate optimal dynamic flux distributions. This entailed uniformly generating random transient concentration profiles within the known experimental constraints, leading to the truncation of the bounded flux polytope—a modality prevalently utilized for characterizing metabolic solution spaces (Herrmann et al., 2019; Schellenberger & Palsson, 2009). To enable an efficient sampling process and mitigate the inefficiencies inherent in rejection sampling, the r-DMFA framework was further modified by incorporating a global optimization strategy facilitated by the BARON global solver (Amaran & Sahinidis, 2012).

Following this, a flux modulation ratio was computed for each culture, defined as the fold changes of each flux in the generated ensemble dynamic model relative to the average representative flux distribution per culture. These flux fold changes served to characterize the influence of each reaction on the variations observed within the ensemble of dynamic flux modulation ratios, providing insight into the reaction-driven flux alterations across different cultures.

#### Generating perturbation models by knockouts (KO)

Perturbations were induced through single enzyme and reaction knockouts, performed by leveraging additional constraints to the r-DMFA framework, grounded on the Minimization of Metabolic Adjustment (MOMA) principle (Segrè et al., 2002), to predict dynamic model responses. In other words, our MOMA-based r-DMFA framework employed quadratic programming, relaxing optimal wild-type flux assumptions, facilitating precise metabolic phenotype predictions of mutants, and elucidating cellular adaptation to enzymatic dysfunction. Furthermore, this approach utilized convex transformation methods in the global optimization algorithm of the BARON solver (Nohra & Sahinidis, 2018). Moreover, the perturbations could take on the form of enzyme overexpression, inhibition, or knockout and is formulated as an additional constraint to the problem in equation (5) given by:

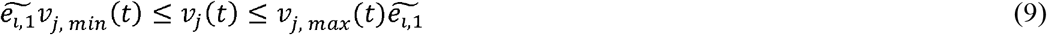

where, 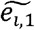, represents the introduced regulation fold change in enzyme expression compared to the reference state.

After each perturbation, the goal was to compare the predicted responses from single knockouts against the experimental wild-type (WT) predictions. An evaluation metric, *z*_*d*_, and spearman’s rank correlation was employed to evaluate the metabolic distance between each mutant dynamic flux phenotype and the WT dynamic flux phenotype. A supplementary module was integrated into the original framework to facilitate these modifications. Additionally, a modified version of the MATLAB implementation as developed by Leighty and Antoniewicz (2011) (Leighty & Antoniewicz, 2011) was utilized to conduct the simulations in this study. The dynamic model along with all requisite scripts for generating the results of this study are made accessible via the link: https://github.com/ssbio/r-DMFA.

### Quantifying metabolic distances in dynamic perturbation models

An evaluation metric, defined as the average relative deviation between wild-type (WT) determined fluxes and those predicted by perturbed (mutant) dynamic models, was employed. The metric utilizes the coefficient of variation (Isaacs et al., 2003) to scale the terms in the metric function, *z*_*d*_, thereby capturing potential degenerate polytopic uncertainty associated with experimental measurements. Consequently, reactions with a tighter confidence interval contribute more significantly to the function. The metric is expressed as:

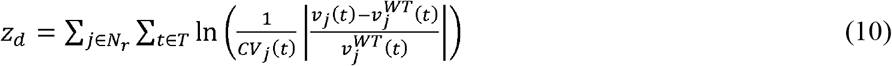

where, 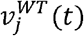 represents the WT flux profile of reaction *j*; *CV*_j_(*t*) denotes the coefficient of variation for each mutant dynamic flux of reaction *j* at each timepoint, employed to assess the relative dispersion of the flux-space solution with sensitivity to experimental measurement uncertainty.*N*_*T*_ represents the number of considered timepoints of *T*, the time domain [*t*_*1*,…,_*t*_*n*_]; *Nr* is the set of nonzero flux reactions in the WT dynamic model. It is noteworthy that reactions that do not carry flux for the WT may exhibit nonzero flux in a perturbed dynamic model. Excluding zero-flux WT reactions from the comparative measure of metabolic distance enables a rigorous and relative comparison across mutants. Mathematically, this excludes jump discontinuities.

Additionally, to assess the metabolic differences between mutant and wild-type (WT) strains, Spearman’s rank correlation was utilized. This method calculates Spearman’s rank correlation coefficients for each metabolic reaction across various time points. Accompanying each output correlation coefficient, designated as rho, is a statistical measure, the p-value, indicating statistical significance. From this, two primary metrics were derived: the ratios of positive correlation reactions and negative correlation reactions. These metrics represent the proportion of reactions displaying significant positive (rho > 0.5, p < 0.05) and negative (rho < −0.5, p < 0.05) correlations, respectively. This provides a succinct depiction of metabolic congruence or divergence between the two strains.

### Statistical analyses and regression analyses

Within this study, various statistical and analytical tools were employed. The two-sample Kolmogorov-Smirnov test (KS test) (Frank J. Massey, 1951) was utilized to compare distributions, while the Bhattacharya coefficient was employed for overlap measurement between distributions. Bootstrap resampling was employed to provide estimation accuracy, and linear regression was used for examining relationships between variables. Additionally, Spearman’s rank correlation coefficient was implemented to evaluate the monotonic relationships between dynamic flux distributions, particularly given that the data in this study did not satisfy the requirements of linearity and homoscedasticity, thereby providing a non-parametric alternative to assess associations without assuming a specific distribution.

## Acknowledgement

AO wishes to express gratitude for the financial support provided for this research. This includes the NIH MIRA Award (5R35GM143009) awarded to RS and the National Science Foundation grant (MCB 1818297) awarded to EBC and RS. We gratefully acknowledge the source code to the initial L-DMFA framework provided by Maciek Antoniewicz from University of Michigan.

## Author contributions

All authors designed research, wrote, and edited the manuscript.

