## Supporting Information for "Quantitative Dynamic Analysis of *de novo* Sphingolipid Biosynthesis in *Arabidopsis thaliana*"

### Supplementary Information

Supplementary Fig. 1:

Growth profiles for the experimental ^15^N labeled and the ^14^N control labeled cultures.

mg dry weight

| Time (days) | 14N | 15N |
| --- | --- | --- |
| 0 | 9.6 | 6.2 |
| 1 | 10.7667 | 19.5 |
| 2 | 19.575 | 18.95 |
| 3 | 23.775 | 20.85 |
| 4 | 22.275 | 28.9 |
| 5 | 36.725 | 39.8 |
| 6 | 43.675 | 57.75 |
| 7 | 53.85 | 63.85 |
| 8 | 58 | 55.45 |
| 9 | 44.425 | 52.1 |
| 12 | 37.6 | 43.9 |
| 15 | 40.625 | 40 |

Supplementary Fig. 2: Temporal abundance of (^14^N + ^15^N) in sphingolipid species highlighting the *de novo* synthesis.

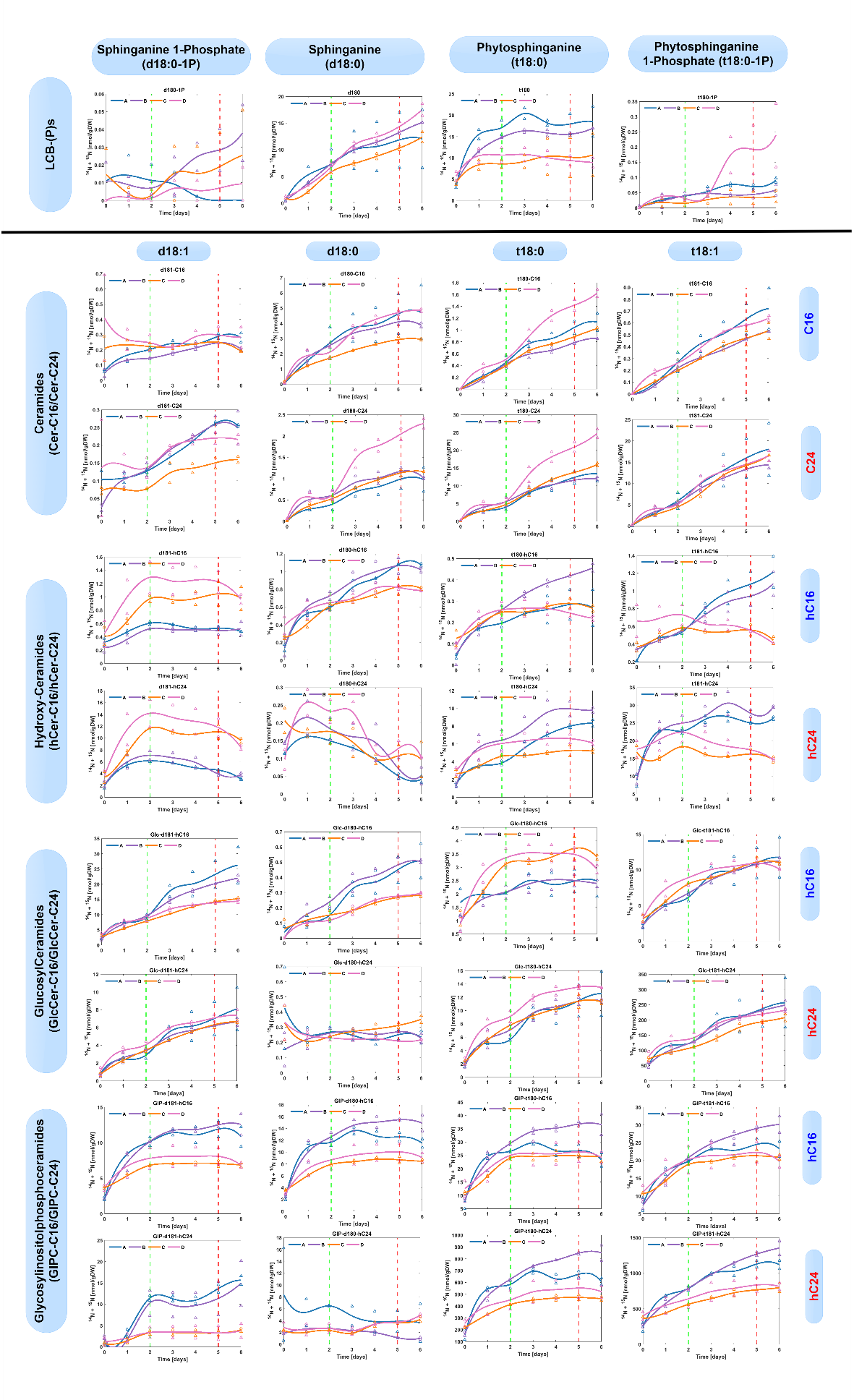

Supplementary Fig. 3: **Temporal abundance of ^15^N in sphingolipid species highlighting the *de novo* synthesis**. This figure showcases the corresponding temporal ^15^N abundances for the 36 sphingolipid species, to emphasize the de-novo synthesis rate across biological replicates, A, B, C and D. Cultures C and D are ^15^N labeled replicates, while A and B are ^14^N unlabeled replicates. The ^15^N enrichment reaches a plateau of 80-100% by the mid-exponential growth phase (day 5), indicating the heightened rate of new sphingolipid synthesis during this phase. The data also present a comparative analysis of the “heavy” (m + 1) isotopologue fraction relative to the unlabeled condition, offering insight into the efficacy of nitrogen isotopes for dynamic labeling purposes.

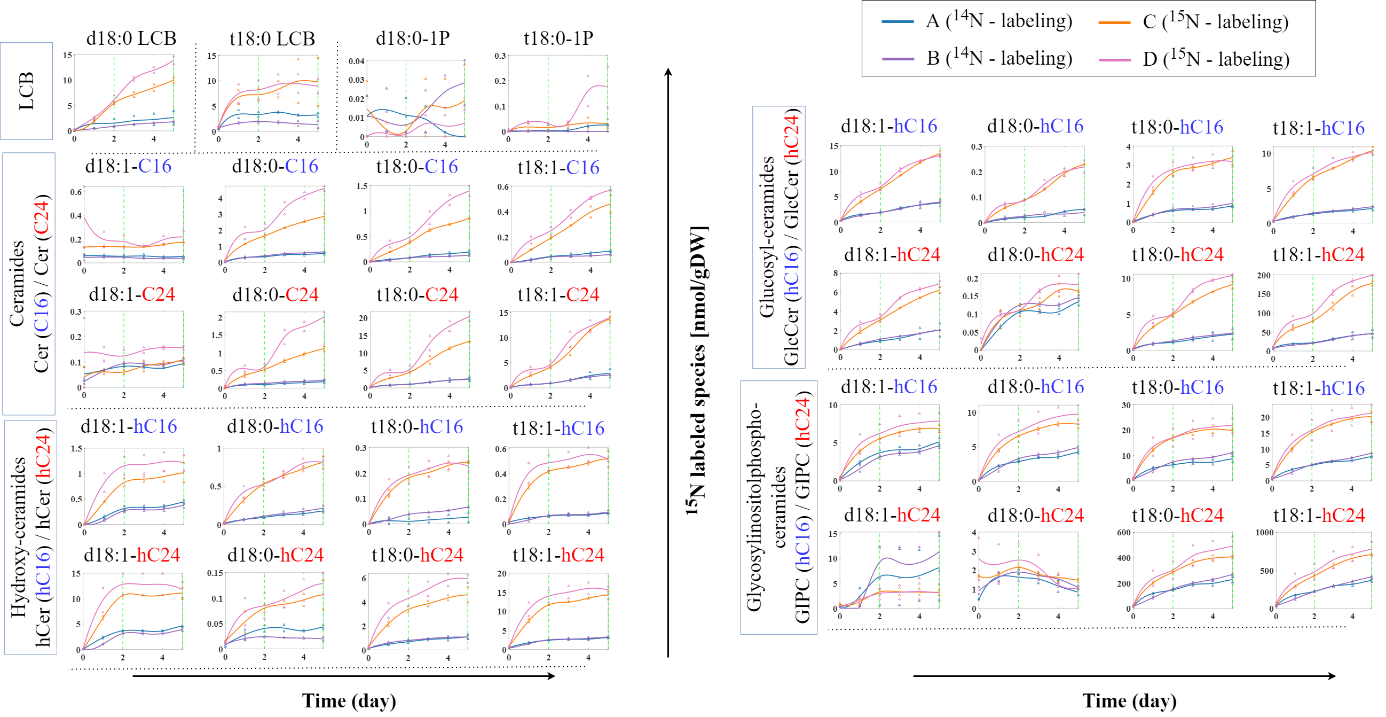

Supplementary Table 1:

Model description describing the blueprint pathway for *de novo* sphingolipid synthesis consisting metabolic transformations annotated by Alsiyabi et al. (2021).

| **Enzyme** | **Reaction ID** | **Reaction Name** | **GPR (gene symbol)** | **Reaction Equation** |
| --- | --- | --- | --- | --- |
| SPT | SPT | Serine palmitoyltransferase | (LCB1 and (LCB2a or LCB2b) and (ssSPTa or ssSPTb)) | Ser => 3KS |
| KSR | KSR | 3-ketosphinganine reductase | (KSR1 or KSR2) | 3KS => d180 |
| SBH | SBH | sphingoid-base hydroxylase | (SBH1 or SBH2) | d180 => t180 |
| LCBK | LCBKa | LCB kinase | (AtLCBK1 or SPHK1 or SPHK2) | d180 => d180-1P |
| LCBK | LCBKb | LCB kinase | (AtLCBK1 or SPHK1 or SPHK2) | t180 => t180-1P |
| LOH2 | CS1a | class I ceramide synthase | LOH2 | d180 => d180-C16 |
| LOH2 | CS1b | class I ceramide synthase | LOH2 | t180 => t180-C16 |
| LOH1/3 | CS2a | class II ceramide synthase | (LOH1 or LOH3) | d180 => d180-C24 |
| LOH1/3 | CS2b | class II ceramide synthase | (LOH1 or LOH3) | t180 => t180-C24 |
| SLD | SLDa | sphingolipid ∆8-desaturase | (SLD1 or SLD2) | d180-C16 => d181-C16 |
| SLD | SLDb | sphingolipid ∆8-desaturase | (SLD1 or SLD2) | t180-C16 => t181-C16 |
| SLD | SLDc | sphingolipid ∆8-desaturase | (SLD1 or SLD2) | d180-C24 => d181-C24 |
| SLD | SLDd | sphingolipid ∆8-desaturase | (SLD1 or SLD2) | t180-C24 => t181-C24 |
| FA2H | FA2Ha | fatty acid 2-hydroxylase | FAH2 | d180-C16 => d180-hC16 |
| FA2H | FA2Hb | fatty acid 2-hydroxylase | FAH2 | t180-C16 => t180-hC16 |
| FA2H | FA2Hc | fatty acid 2-hydroxylase | FAH1 | d180-C24 => d180-hC24 |
| FA2H | FA2Hd | fatty acid 2-hydroxylase | FAH1 | t180-C24 => t180-hC24 |
| FA2H | FA2He | fatty acid 2-hydroxylase | FAH2 | d181-C16 => d181-hC16 |
| FA2H | FA2Hf | fatty acid 2-hydroxylase | FAH2 | t181-C16 => t181-hC16 |
| FA2H | FA2Hg | fatty acid 2-hydroxylase | FAH1 | d181-C24 => d181-hC24 |
| FA2H | FA2Hh | fatty acid 2-hydroxylase | FAH1 | t181-C24 => t181-hC24 |
| GCS | GCSa | glucosylceramide synthase | GCS | d180-hC16 => Glc-d180-hC16 |
| GCS | GCSb | glucosylceramide synthase | GCS | t180-hC16 => Glc-t180-hC16 |
| GCS | GCSc | glucosylceramide synthase | GCS | d180-hC24 => Glc-d180-hC24 |
| GCS | GCSd | glucosylceramide synthase | GCS | t180-hC24 => Glc-t180-hC24 |
| GCS | GCSe | glucosylceramide synthase | GCS | d181-hC16 => Glc-d181-hC16 |
| GCS | GCSf | glucosylceramide synthase | GCS | t181-hC16 => Glc-t181-hC16 |
| GCS | GCSg | glucosylceramide synthase | GCS | d181-hC24 => Glc-d181-hC24 |
| GCS | GCSh | glucosylceramide synthase | GCS | t181-hC24 => Glc-t181-hC24 |
| GIPCS | GIPCSa | GIPC synthase | ((IPCS1 or IPCS2 or IPCS3) and IPUT1) | d180-hC16 => GIP-d180-hC16 |
| GIPCS | GIPCSb | GIPC synthase | ((IPCS1 or IPCS2 or IPCS3) and IPUT1) | t180-hC16 => GIP-t180-hC16 |
| GIPCS | GIPCSc | GIPC synthase | ((IPCS1 or IPCS2 or IPCS3) and IPUT1) | d180-hC24 => GIP-d180-hC24 |
| GIPCS | GIPCSd | GIPC synthase | ((IPCS1 or IPCS2 or IPCS3) and IPUT1) | t180-hC24 => GIP-t180-hC24 |
| GIPCS | GIPCSe | GIPC synthase | ((IPCS1 or IPCS2 or IPCS3) and IPUT1) | d181-hC16 => GIP-d181-hC16 |
| GIPCS | GIPCSf | GIPC synthase | ((IPCS1 or IPCS2 or IPCS3) and IPUT1) | t181-hC16 => GIP-t181-hC16 |
| GIPCS | GIPCSg | GIPC synthase | ((IPCS1 or IPCS2 or IPCS3) and IPUT1) | d181-hC24 => GIP-d181-hC24 |
| GIPCS | GIPCSh | GIPC synthase | ((IPCS1 or IPCS2 or IPCS3) and IPUT1) | t181-hC24 => GIP-t181-hC24 |
| DPL1 | DPL1a | dihydrosphingosine phosphate lyase | AtDPL1 | d180-1P => C16O |
| DPL1 | DPL1b | dihydrosphingosine phosphate lyase | AtDPL1 | t180-1P => C16O |

Model Compounds:

| **ID** | **Name** |
| --- | --- |
| Ser | Serine |
| 3KS | 3-ketosphinganine |
| d180 | Sphinganine |
| t180 | Phytosphingosine |
| d180-1P | Sphinganine 1-Phosphate |
| t180-1P | Phytosphinganine 1-Phosphate |
| d180-C16 | Dihydroceramide (C16 FA) |
| t180-C16 | Phyto(hihydro)ceramide (C16 FA) |
| d180-C24 | Dihydroceramide (C24 FA) |
| t180-C24 | Phyto(hihydro)ceramide (C24 FA) |
| d181-C16 | ∆8 unsat. Dihydroceramide (C16 FA) |
| t181-C16 | ∆8 unsat. Phyto(hihydro)ceramide (C16 FA) |
| d181-C24 | ∆8 unsat. Dihydroceramide (C24 FA) |
| t181-C24 | ∆8 unsat. Phyto(hihydro)ceramide (C24 FA) |
| d180-hC16 | hydroxy-d180-C16 |
| t180-hC16 | hydroxy-t180-C16 |
| d180-hC24 | hydroxy-d180-C24 |
| t180-hC24 | hydroxy-t180-C24 |
| d181-hC16 | hydroxy-d181-C16 |
| t181-hC16 | hydroxy-t181-C16 |
| d181-hC24 | hydroxy-d181-C24 |
| t181-hC24 | hydroxy-t181-C24 |
| Glc-d180-hC16 | d180-hC16 GlcCer |
| Glc-t180-hC16 | t180-hC16 GlcCer |
| Glc-d180-hC24 | d180-hC24 GlcCer |
| Glc-t180-hC24 | t180-hC24 GlcCer |
| Glc-d181-hC16 | d181-hC16 GlcCer |
| Glc-t181-hC16 | t181-hC16 GlcCer |
| Glc-d181-hC24 | d181-hC24 GlcCer |
| Glc-t181-hC24 | t181-hC24 GlcCer |
| GIP-d180-hC16 | d180-hC16 GIPC |
| GIP-t180-hC16 | t180-hC16 GIPC |
| GIP-d180-hC24 | d180-hC24 GIPC |
| GIP-t180-hC24 | t180-hC24 GIPC |
| GIP-d181-hC16 | d181-hC16 GIPC |
| GIP-t181-hC16 | t181-hC16 GIPC |
| GIP-d181-hC24 | d181-hC24 GIPC |
| GIP-t181-hC24 | t181-hC24 GIPC |
| C16O | hexadec-6-ene-1,2-diol |
